# MINE is a method for detecting spatial density of regulatory chromatin interactions based on a MultI-modal NEtwork

**DOI:** 10.1101/2022.07.11.499656

**Authors:** Haiyan Gong, Minghong Li, Mengdie Ji, Xiaotong Zhang, Zan Yuan, Sichen Zhang, Yi Yang, Chun Li, Yang Chen

## Abstract

Chromatin interactions play essential roles in chromatin conformation and gene expression. However, few tools exist to analyze the spatial density of regulatory chromatin interactions. Here, we present the MultI-modal NEtwork (MINE) toolkit, including MINE-Loop, MINE-Density, and MINE-Viewer. MINE-Loop network modeling integrates Hi-C, ATAC-seq, and histone ChIP-seq data to enhance the detection of regulatory chromatin interactions (RCIs, *i*.*e*., chromatin interactions that are anchoring regulatory elements to chromatin); MINE-Density quantifies the spatial density of regulatory chromatin interactions identified by MINE-Loop within different chromatin conformations; and MINE-Viewer facilitates 3D visualization of the density of chromatin interactions and participating regulatory factors, such as transcription factors. We applied MINE to investigate the relationship between the spatial density of regulatory chromatin interactions (SD-RCI) and chromatin volume change in HeLa cells before and after liquid-liquid phase separation. Changes in SD-RCI before and after treating the HeLa cells with 1,6-hexanediol showed that the change in chromatin volume was related to the degree of activation or repression of genes in chromatin interactions. The MINE toolkit provides a new method for quantitative study of chromatin conformation.

**Graphical Abstract:** 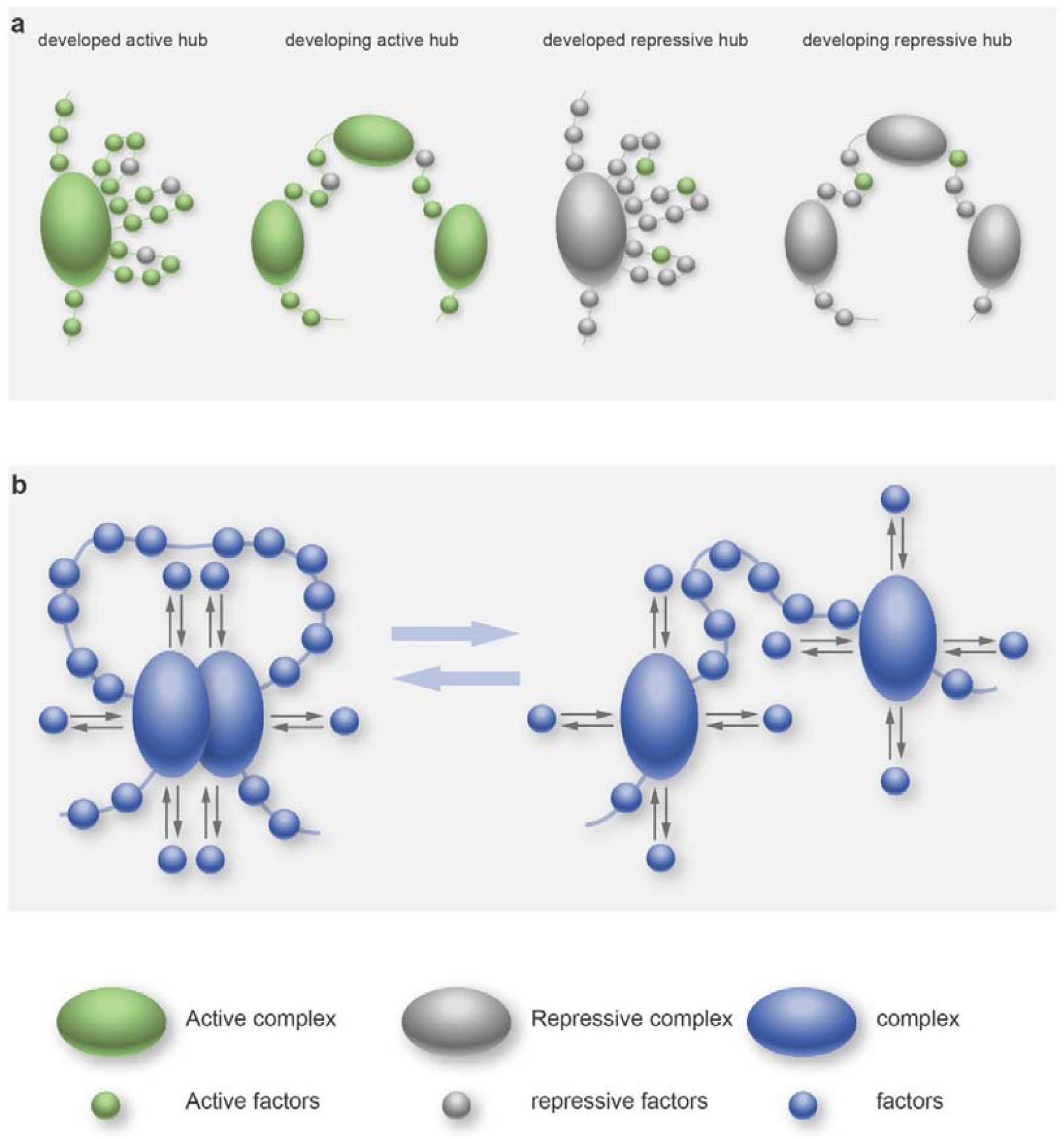

**In Brief:** Gong et al. provide a toolkit MINE to explore the relationship between spatial density of regulatory chromatin interactions, gene expression and chromatin structure change.

**Highlights:**

- MINE toolkit was provided to detect more regulatory chromatin interactions (RCI), count the spatial density of regulatory chromatin interactions and visualize the density of chromatin interactions with transcription factors in three-dimension.
- Chromatin regions were divided into developed active hub, developing active hub, developed repressive hub, and developing repressive hub according to the density of active or repressive regulatory chromatin interactions.
- The change of chromatin structure before and after liquid-liquid phase separation is quantitively described by using the MINE toolkit.

## INTRODUCTION

With advances in 3D genome research, increasing evidence supports an essential role for chromatin interactions with nuclear regulatory factors in shaping chromatin conformation and the regulation of gene transcription. For example, the chromatin structure of A/B compartments and topologically associating domains (TADs) appear to be formed through distinct chromatin interactions (*i*.*e*., loops). (Wit, 2019) Previous research has revealed that gene densities and GC content are correlated with the density of chromatin interactions (*i*.*e*., the number of interactions per Mb). (Sandhu et al., 2012) Hou et al. showed that gene density and transcription contribute to the partition of physical domains (*i*.*e*., regions with high gene density) (Hou et al., 2012). Interactions between chromatin features associated with transcriptional activation or repression, such as Rad21, CTCF, and H3K4me3, are correlated with gene expression (Seitan et al., 2013). L. Almassalha et al. provided a “macrogenomic engineering” approach to regulate transcriptional activity in cancer cells by modulating chromatin density (Almassalha et al., 2017). Therefore, research on the spatial density of regulatory chromatin interactions can further our understanding of the mechanisms responsible for chromatin folding and the relationship between chromatin conformation and gene expression. ChIA-PET (Fullwood and Ruan, 2009) and HiChIP (Mumbach et al., 2016) are tools that can capture the regulatory chromatin interactions for proteins of interest, such as DNA-binding regulatory proteins, RNA transcription factors. However, ChIA-PET and HiChIP data are only available for some cell lines, and their acquisition for different target proteins is costly, laborious, and time-consuming. Hi-C (Van Berkum et al., 2010) is a sequencing technology used to quantify the number of interactions between genome bins adjacent in 3D space but may farther in a linear genome. Therefore, a method to detect regulatory chromatin interactions by calling loops from Hi-C data can help reduce the cost. In this paper, we proposed MINE-Loop to identify special regulatory chromatin interactions from high-resolution Hi-C data.

Among the numerous physical mechanisms of chromatin formation, one type of model (Brackley et al., 2016; Di Pierro et al., 2016; Di Stefano et al., 2016; Fiorillo et al., 2020) considers the formation of chromatin structures mediated by chromatin interactions with molecular factors, such as architectural proteins, histone marks and non-coding RNAs (Jung and Kim, 2021). Specifically, the Strings and Binders Switch (SBS) model proposes that chromatin is a ‘self-avoiding polymer’ surrounded by diffusive molecular factors (*e*.*g*., transcription factors) that anchor to cognate recognition sites on the chromosome to drive the chromatin folding process. The SBS model can be specifically applied to study the relationship between chromatin structural states, such as loops, TADs, or A/B compartments, and the density of regulatory factors (*i*.*e*., regulatory elements, such as enhancers, promoter and silencer). Studies investigating loops have shown that CCCTC-Binding Factor (CTCF) mediates interactions with chromatin regions enriched with enhancer-regulated genes by altering chromatin domain structures (Oti et al., 2016; Ren et al., 2017). Similarly, Golkaram et al. (Golkaram et al., 2017) examined local chromatin density to quantify transcriptional regulatory components in a cell population and found that distinct topologically associating domains (TADs) determined the distribution of gene expression. In a study of open chromatin domains, Jiang et al. proposed the spatial density of open chromatin metric to characterize intra-TAD chromatin state and structure, and found that TADs with decreased SDOC were enriched with repressed genes during T-cell development in mice (Jiang et al., 2021). While these studies investigated the relationship between chromatin structure and specific molecular factors (*e*.*g*., transcription factors), a method for simultaneous quantification of the spatial density of active or repressive regulatory chromatin interactions (RCIs, *i*.*e*., chromatin interactions that are anchoring regulatory elements to chromatin) is still lacking. Such a method could provide quantitative evidence supporting or refuting the SBS model of the relationship between chromatin structure and gene expression.

To address this gap, we introduce the MultI-modal NEtwork (MINE) data analysis toolkit, including the MINE-Loop, MINE-density, and MINE-Viewer tools, to explore the spatial density of regulatory chromatin interactions. MINE-Loop is a neural network model that integrates Hi-C, ChIP-seq (Schmidt et al., 2009), and ATAC-seq (Buenrostro et al., 2015) data to enhance the proportion of detectable regulatory chromatin interactions by reducing noise from non-regulatory interactions. MINE-Density can be used to calculate the spatial density of regulatory chromatin interactions (SD-RCI) identified by MINE-Loop, and MINE-Viewer facilitates visualization of density and specific interactions with regulatory factors in 3D genomic structures. We explored the relationship between SD-RCI, calculated by MINE-density, and the status of gene transcription in HepG2 cells and identify four distinct levels of interaction density related to the formation of transcriptionally active or repressive chromatin regions enriched for anchored regulatory factors (*i*.*e*., developed hubs) or regions with low density of anchored factors (developing hubs). Finally, we applied SD-RCI values to data obtained in a liquid-liquid phase separation experiment (*i*.*e*., the HeLa cell line treated or not with 1,6-hexanediol) to quantitatively describe volume changes in chromosome structure, which revealed that nucleus expands after treating with 1,6-hexanediol. In summary, MINE provides a new method for quantitative analysis of chromatin conformations.

## RESULTS

### Overview of the MINE framework

MINE is a multi-modal method for detecting the spatial density of regulatory chromatin interactions (RCIs, *i*.*e*., chromatin interactions that are anchoring regulatory elements to chromatin, *e*.*g*., CTCF, RAD21, SMC3, POLR2A) that includes MINE-Loop, MINE-Density, and MINE-Viewer functions (Figure 1A (left)).

**Figure 1.**
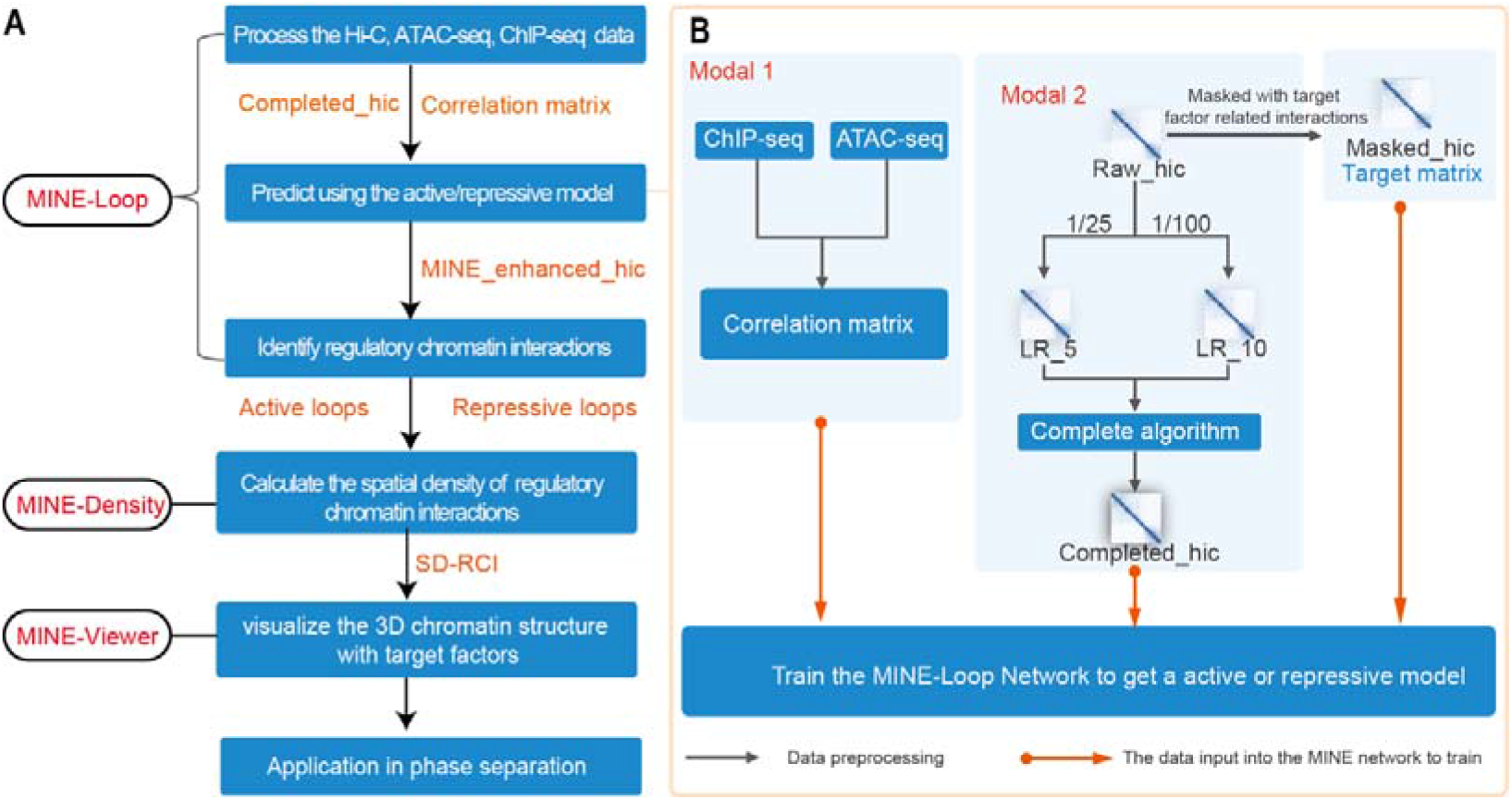
overview of the MINE pipeline. (A) Workflow and analytical pipeline of the MINE method. (B) An overview of the MINE-Loop architecture and workflow for model training.

#### (1) Description of the MINE-Loop tool

Since raw Hi-C deep sequencing data contains a high degree of noise, currently available loop (*i*.*e*., Two chromatin regions that the interaction frequency is higher than that of the surrounding adjacent regions in the Hi-C contact matrix.) callers commonly identify a relatively low proportion of RCIs. Hence, MINE-Loop was developed to obtain a larger proportion of RCIs from enhanced Hi-C data than from raw Hi-C data.

As shown in Figure 1B (right), the MINE-Loop tool is first generating a high-resolution (*i*.*e*., 1 kb resolution) VC-normalized Hi-C contact matrix (Raw_hic) from raw Hi-C deep sequencing data obtained from human GM12878, H1-hESC, and HepG2 cell lines (build GRCh38). We then generated a masked Hi-C matrix (Masked_hic) from the Raw_hic file using ATAC-seq or histone ChIP-seq data obtained from the same cell line, resulting in a targeted Hi-C matrix that contains a large proportion of regulatory chromatin interactions that are used to train the MINE-Loop network.

Since the peaks called from the ATAC-seq and histone ChIP-seq data can indicate the localization of regulatory chromatin interactions, we next generated a correlation matrix of ATAC-seq and ChIP-seq peaks by calculating the Pearson Correlation Coefficients between the peaks of these datasets. This correlation matrix serves as the first modal matrix for training the MINE-Loop model.

We then processed the Raw_hic matrix to generate the second modal matrix of training the MINE-Loop model. The Raw_hic matrix was down-sampled at a ratio of 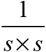 to build a smoothed Hi-C contact matrix at the same resolution as Raw_hic, where the values of the *s* × *s* window were set as the average values of the *s* × *s* window. The essential reason to smooth Raw_hic matrix is that the exact location of loops cannot be determined. We think this is beneficial for reducing noise. To extract more features of the Hi-C matrix, we then processed these down-sampled Hi-C matrices by point-to-point addition, and enhanced the Hi-C matrix after addition using the FAN method (Achanta et al., 2017) to obtain a completed Hi-C matrix (Completed_hic).

Finally, the correlation matrix, Completed_hic and Masked_hic are fed into the MINE-Loop network to map functions among the correlation matrix, Completed_hic and Masked_hic. Once the model is trained, it is then applied to generate an enhanced Hi-C contact matrix (MINE_enhanced_hic) for any cell line using the Completed_hic and correlation matrix as inputs. To identify RCIs, MINE_enhanced_hic can then be fed into two available loop calling tools (*i*.*e*., FitHiC2 (Kaul et al., 2020) and mustache (Ardakany et al., 2020)).

The MINE-Loop model can increase the proportion of active or repressive RCIs by training with ChIP-seq of different histone modifications. For ChIP-seq data of different active-related histone modifications (*i*.*e*., H3K27ac, H3K4me3), an active MINE-Loop model (*i*.*e*., active model) can obtain a larger proportion of RCIs related to the control of DNA transcription machinery, while ChIP-seq data of suppression-related histone modifications (*i*.*e*., H3K27me3, H3k9me3) can be used to train repressive MINE-Loop model (*i*.*e*., repressive model) to obtain a larger proportion of transcriptionally repressive chromatin interactions. We define loops called from active model as active loops, loops called from repressive model as repressive loops.

#### (2) Description of the MINE-density and MINE-Viewer tools

Based on the RCIs identified by the MINE-Loop network, the ratio of the total number of active or repressive loops in a TAD to the entire 3D space physically occupied by the TAD structure is defined as the spatial density of RCIs (SD-RCI). The 3D chromatin structure can be visualized with target factors (*e*.*g*., CTCF, genes, H3K4me3 and POLR2A) using the MINE-Viewer tool to investigate the density of target factor distribution in the 3D chromatin structure. Ultimately, MINE is applied to analyze changes in the spatial density of RCIs before and after phase separation.

### MINE-Loop facilitates detecting a high proportion of regulatory chromatin interactions

#### (1) MINE-Loop can detect regulatory chromatin interactions in high-noise data

We next assessed whether the MINE-Loop analysis could increase the proportion of detectable RCIs compared with that obtained by current loop callers from Raw_hic using an active model as an example. We first generated the Completed_hic, correlation matrix and Masked_hic of the GM12878 cell line that targeted factors (*e*.*g*., H3K4me3 and H3K27ac) specifically involved in DNA transcription to train and test the active model.

Hi-C data from the GM12878 cell line was downloaded from the 4DNucleome database (https://data.4dnucleome.org/ (Dekker et al., 2017); accession number 4DNFI1UEG1HD (Rao et al., 2014)) to generate the Completed_hic. The ATAC-seq and H3K27ac, H3K4me3 histone ChIP-seq data from the GM12878 cell line was downloaded from the ENCODE database (accession numbers ENCSR637XSC (Consortium, 2012), ENCSR057BWO (Lee et al., 2020), and ENCSR000AKC (Lee et al., 2020)) and subsequently used to generate the correlation matrix. The annotation file of candidate Cis-Regulatory Elements (CREs) in GM12878 cells was downloaded from ENCODE (accession number ENCSR820WFY (Moore et al., 2020)) to generate the Masked_hic.

Then, matrices generated for human chromosomes 1-17 in GM12878 cell line dataset were used for training the active model, while the matrices generated for human chromosomes 18-23 in GM12878 dataset were used to test the performance of the active model with the enhanced Hi-C data in the MINE_enhanced_hic output file.

To test the active model, FitHiC2 (Kaul et al., 2020) and mustache (Ardakany et al., 2020) were used to call intrachromosomal loops within a genomic distance of 2-100 kb in the MINE_enhanced_hic and Raw_hic matrices of human chromosomes 18-23 in GM12878 cells. The results showed a 12.5% and 9.7% overlap between loops called from MINE_enhanced_hic and Raw_hic data by FitHiC2 and mustache (Figure S2, S3 in Supplementary Files 2). However, the loops called from the MINE_enhanced_hic had 3-4 times more transcription factors (TFs) related to the control of DNA transcription machinery (*e*.*g*., CTCF, RAD21, SMC3, POLR2A) anchored at the same corresponding loops and transcription start sites (TSS) than that in Raw_hic data (Figure S2 b-f and Figure S3 b-e in Supplementary Files 2).

The same comparison was conducted for called loops ranging from 2-300 kb and 2-500 kb, which showed a greater number of anchored factors in the enhanced Hi-C data (Figure S2 in Supplementary Files 2). These results suggested that MINE-Loop analysis could also reveal long-range regulatory chromatin interactions.

#### (2) The general applicability of MINE-Loop model

Having generated enhanced, integrative datasets related to regulatory chromatin interactions by training an active model, we then explored the effects of using different histone ChIP-seq data combinations to train active models. To this end, three combinations (Combination (i): ATAC-seq, H3K27ac ChIP-seq, H3K4me3 ChIP-seq. Combination (ii): ATAC-seq, H3K27ac ChIP-seq. Combination (iii): ATAC-seq, H3K4me3 ChIP-seq.) were used as inputs for active model training (Figure 2A-D). Examination of anchor number for the CTCF, RAD21, SMC3, and POLR2A TFs in [2, 100 kb] loops (Figure 2A-D) and [2, 300 kb] loops (Figure S4a-d in Supplementary Files 2) indicated that the combination (ii) revealed fewer TF anchor points than other combinations in active interactions. This suggested that the more active-related histone ChIP-seq data used to train active models, the more active TF anchor chromatin interactions will be detectable.

**Figure 2.**
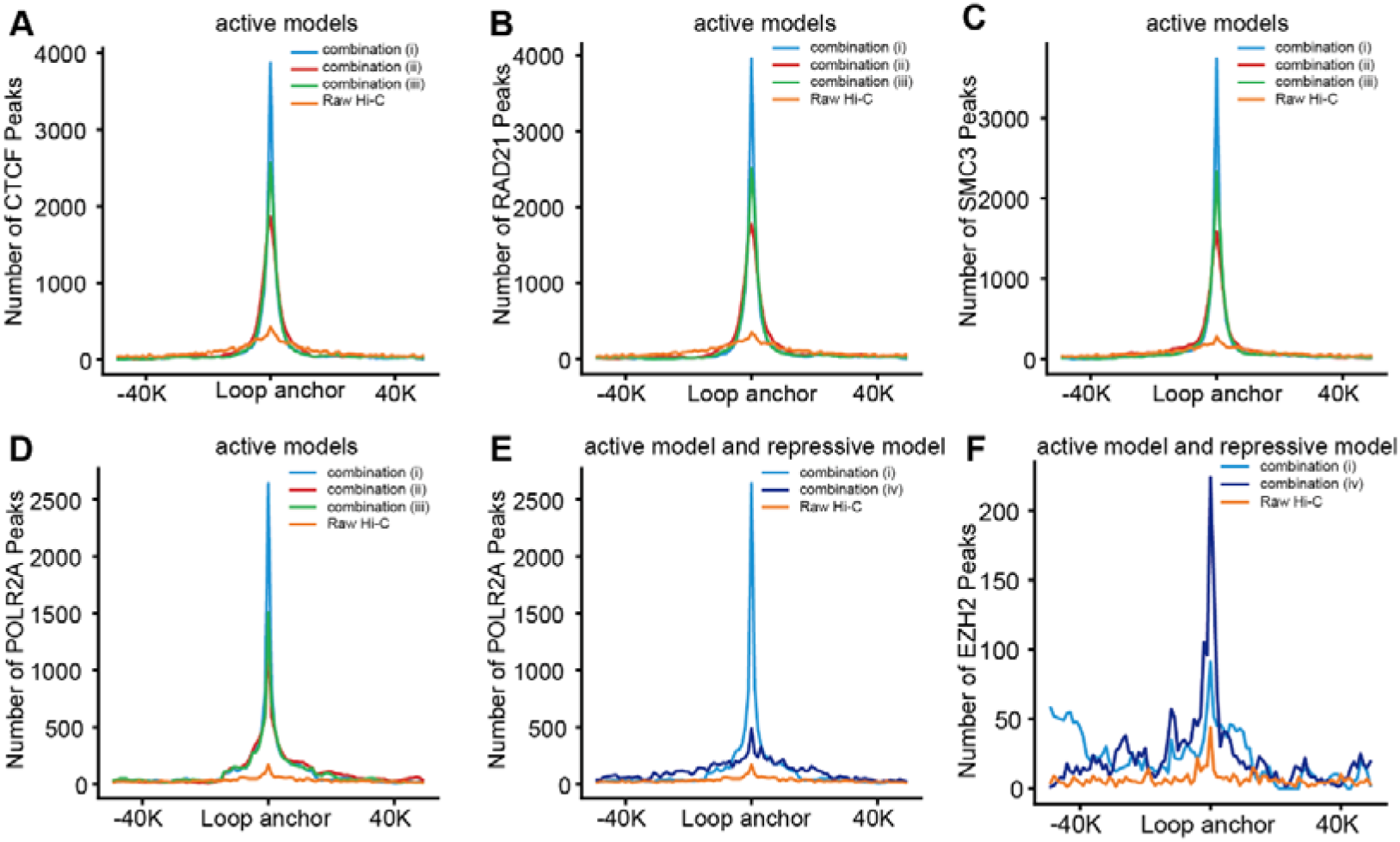
MINE-Loop can detect regulatory chromatin interactions in high-noise data. The active model was trained in the GM12878 cell line using the following three combinations as inputs. Combination (i): ATAC-seq, H3K27ac ChIP-seq, H3K4me3 ChIP-seq (i.e., blue lines). Combination (ii): ATAC-seq, H3K27ac ChIP-seq (i.e., red lines). Combination (iii): ATAC-seq, H3K4me3 ChIP-seq (i.e., green lines). The repressive model was trained using the combination (iv), including H3K27me3 with H3k9me3 ChIP-seq data from the GM12878 cell line (i.e., purple lines). (A–D) Number of CTCF, RAD21, SMC3, POLR2A transcription factor anchors in 2-100kb loops called from active models trained with different combinations of data. (E-F) The comparison of POLR2A and EZH2 anchors in loops called from active and repressive models.

To determine the general applicability of MINE-Loop model, we applied the active model trained using the combination (i) dataset in GM12878 cell line to predict using the combination (i, ii, iii) datasets in IMR90 cell line as input data. The results show that the three combinations all perform better than raw Hi-C data in the number of anchoring TFs (Figure S5 in Supplementary Files 2), suggesting MINE-Loop model do not require all epigenome data that used in the training process. Besides the IMR90 cell line, the data of K562, H1-hESC and HepG2 cell lines is also conducted using the active model trained in GM12878 cell line. The results show that within the genomic distance of 2∼100, loops called from MINE_enhanced_hic in IMR90 (Figure 2G-J) and K562 (Figure S6 in Supplementary Files 2) cell line both can anchor more TFs than from Raw_hic, but less than in GM12878 (Figure 2A-D), H1-hESC (Figure S7 in Supplementary Files 2) and HepG2 (Figure S8 in Supplementary Files 2) cell line. The results suggest that the sequencing depth of the raw Hi-C data has an impact on the effect of MINE-Loop model, where the sequencing depth of K562 and IMR90 cell lines (∼1 billion) is much lower than that of GM12878 (∼4.01 billion), H1-hESC (∼3.22 billion) and HepG2 (2.02 billion) cell lines.

Following the identification of loops containing transcriptional machinery genes that are actively transcribed, we then used a repressive MINE-Loop model (repressive model) trained with suppression-related histone marks (*i*.*e*., H3K27me3, H3K9me3) target ChIP-seq data downloaded from ENCODE (accession number ENCSR000DRX and ENCSR000AOX (Zhang et al., 2020)) to assess whether MINE-Loop could improve detection of transcriptionally repressive chromatin interactions. Comparison of POLR2A and EZH2 (a transcript factor related to long-term transcriptional inhibition) factors between active and repressive models trained with GM12878 cell line data showed that 2∼100 kb loops in the repressive model obtained more EZH2 anchors than those of POLR2A (Figure 2E-F), indicating that active models successfully identified more transcriptional activation-related interactions while the repressive model identified more transcriptional inhibition-related interactions. In agreement with our experimental evaluation of the active model, the repressive model trained with GM12878 cell line data was also used for prediction in other cell lines. In the K562 (Figure S9) and HepG2 (Figure S10) cell lines, loops called from MINE_enhanced_hic revealed a greater number of EZH2 anchors than that obtained from Raw_hic. Collectively, these data demonstrated that the MINE-Loop tool can facilitate detection of a high proportion of active and repressive RCIs.

#### (3) MINE-Loop facilitates the detection of functional chromatin loops

We next sought to verify whether the loops with anchored TFs called from the MINE_enhanced_hic overlapped with ChIA-PET region. We found that 42.43%, 21.64% of the CCCTC-binding factor (CTCF) and POLR2A ChIA-PET region overlapped with the active loops anchored CTCF and POLR2A in HepG2 cell line, suggesting that loops called from MINE_enhanced_hic include many transcription factors binding interactions (Figure S11a-b). The Venn graph of Raw_hic, MINE_enhanced_hic and POLR2A ChIA-PET in GM12878 cell line, showed that MINE_enhanced_hic could overlap with more POLR2A ChIA-PET region (about 9606) than Raw_hic (Figure 3A). The number of anchoring TSS sites around the active loops (Figure 3B) further proved that MINE_enhanced_hic could detect more functional chromatin loops than Raw_hic.

**Figure 3.**
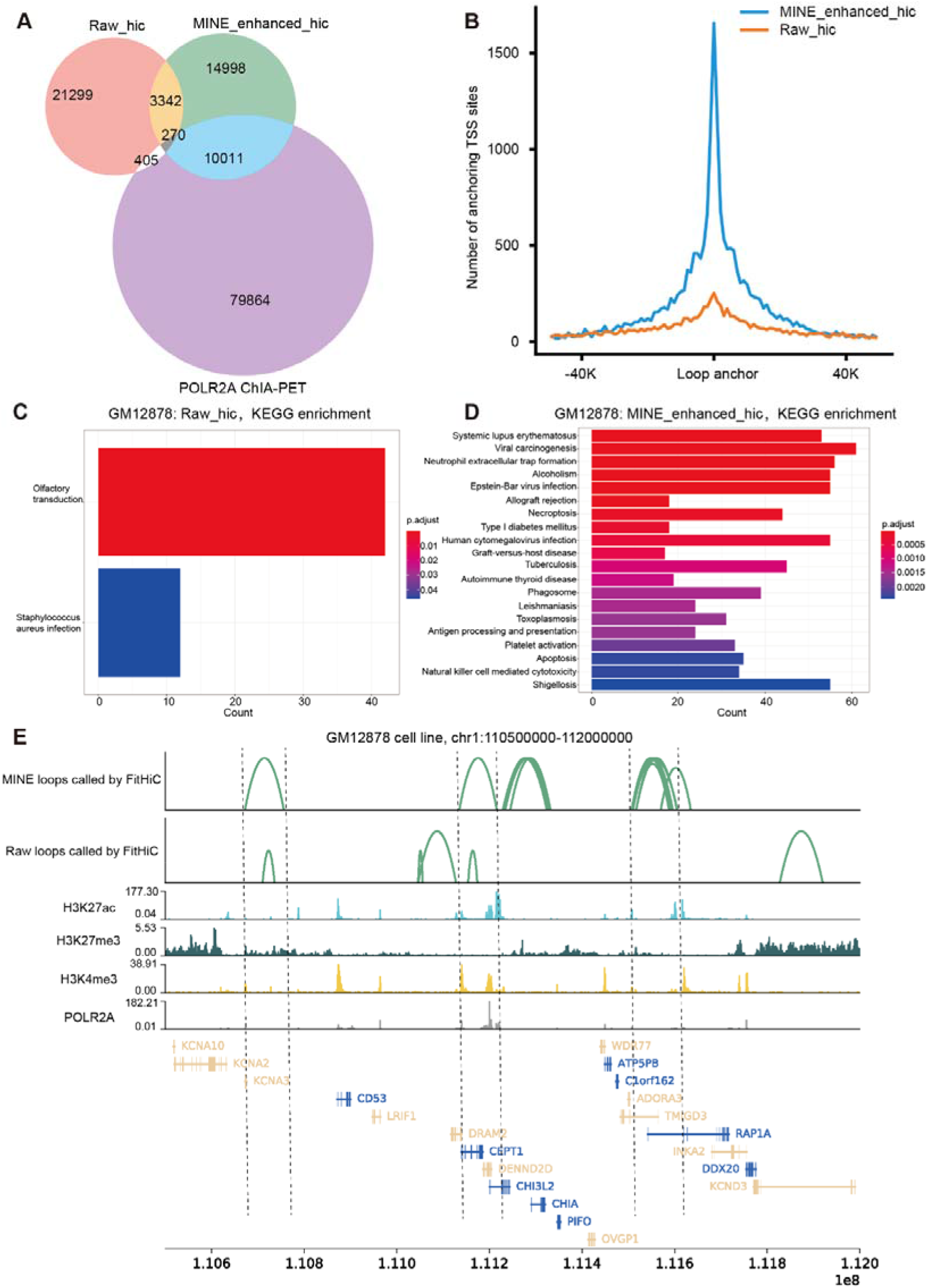
MINE-Loop facilitates the detection of functional chromatin loops. The active model was trained by GM12878 cell line with data of combination (i). (A) The Venn graph of loops called from Raw_hic, MINE_enhanced_hic and POLR2A ChIA-PET data. (B) MINE_enhanced_hic anchor more TSS sites than Raw_hic. (C-D) The differential gene KEGG enrichment of Raw_hic and MINE_enhanced_hic. (E) Visualization of loops, H3K27ac, H3K27me3, H3K4me3, POLR2A target ChIP-seq tracks.

We next performed KEGG enrichment analysis to investigate the predicted functions of genes in active loops that were differentially detected between Raw_hic and MINE_enhanced_hic datasets generated with GM12878 or HepG2 cell lines. The differential genes enriched in active loops called from MINE_enhanced_hic in GM12878 data were involved in ‘Immune-related’ processes (Figure 3C-D). In the HepG2 cell line, the differential genes from MINE_enhanced_hic were enriched in terms related to ‘liver disease’, such as non-alcoholic fatty liver disease, which was consistent with the characteristics of HepG2 cells, while differential genes from Raw_hic were enriched in terms related to ‘sensory perception of smell’, ‘detection of chemical stimulus involved in sensory perception of smell’ (Figure S12a-b). This shows that the differential loops of MINE_enhanced_hic compared to Raw_hic are enriched for genes functionally related to characteristics of that cell line.

To further investigate the enrichment pattern in a locus that loops called from Raw_hic and MINE_enhanced_hic are quite different, we chose the chr1: 110500000-112000000 region in chromosome 1. Upon close inspection of the region (Figure 3E), we found that the differential loops of MINE_enhanced_hic were enriched with H3K27ac, H3K27me3, H3K9me3 and POLR2A signals. These results showed that MINE-Loop could enhance the detection of functional regulatory chromatin interactions.

### Spatial density of regulatory chromatin interactions and gene transcription

To investigate the active and repressive model how to affect the spatial density of regulatory chromatin interactions, we refer to the definition of spatial density of open chromatin (SDOC) metric (Jiang et al., 2021) used to quantitatively measure the intra-TAD chromatin state and structure, and propose the spatial density of regulatory chromatin interactions (SD-RCI) (Figure 4A). The SD-RCI is defined as the ratio of the total number of active or repressive loops in a TAD to the entire 3D space taken up by the physical structure of the TAD. Based on the definition of SD-RCI, the MINE-density tool is developed to calculate the SD-RCI.

**Figure 4.**
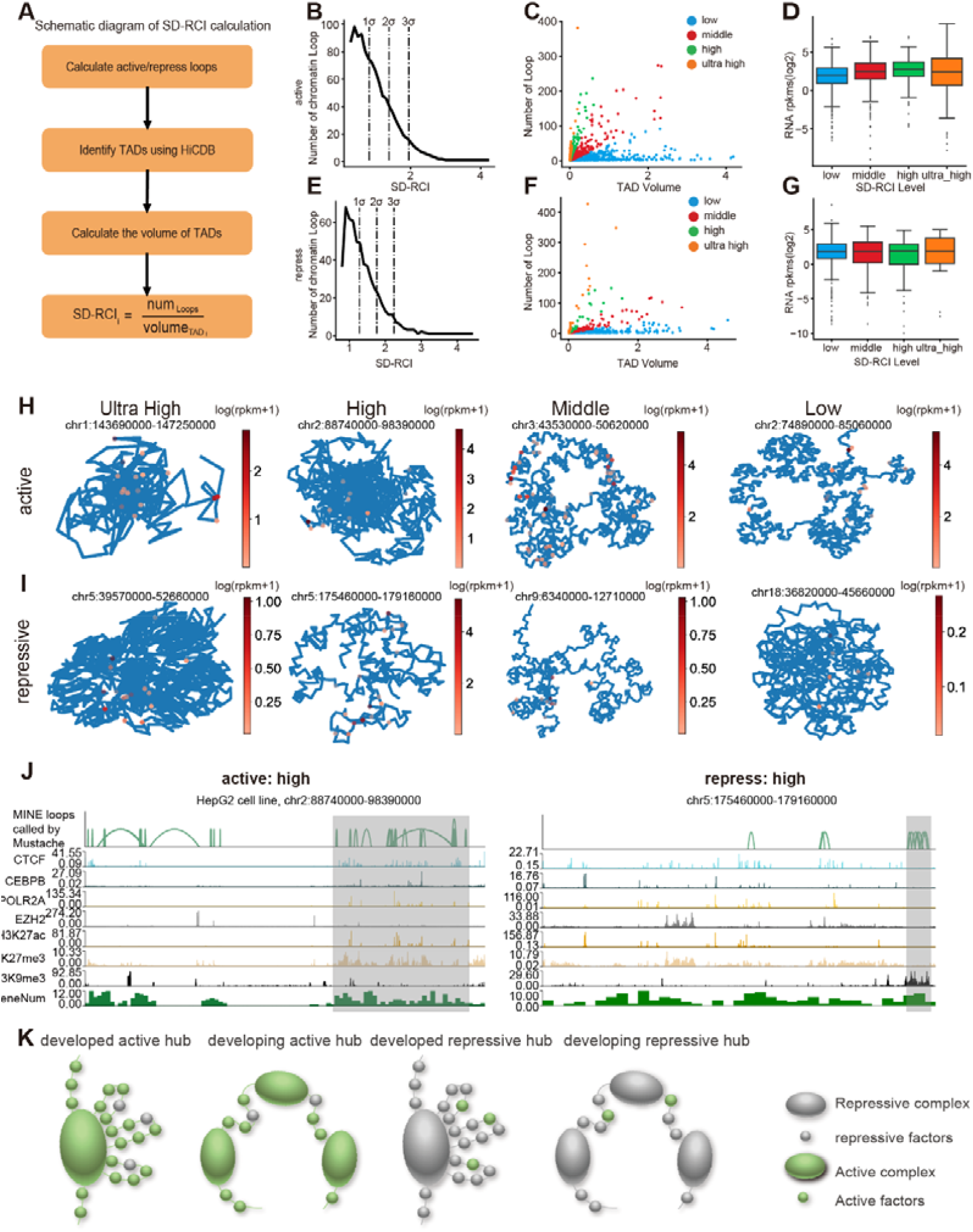
SD-RCI is correlated with gene expression. (A) Schematic diagram of SD-RCI calculation. (B – E) The distribution of the number of active loops changes with the SD-RCI value calculated by the SD-RCI method. (C – F) The dot plot of the number of RCIs and volume of TADs. (D – G) The box plot of RPKM and SD-RCI degree. the SD-RCI is divided into four degrees according to the value of d in the Gaussian distribution of Figure 4A. (H – I) The 3D genome TAD structure visualization with the gene expression strength in four levels from the active and repress model. (J) visualization of loops, CTCF, and histone mark ChIP-seq tracks. Loops are identified from MINE_enhanced_hic by using MUSTACHE. The 3D structure of this region corresponds to the active (repress) high level in Figure 4.(K) Four types of hubs defined by the active or repressive SD-RCI.

To explore the relationship between the SD-RCI and the status of gene transcription in HepG2 cell line, we first generated active and repressive loops within a genomic distance of 2-100 kb (Figure 4) and 2-300 kb (Figure S13d-f) using the active model and repress model. Based on the active and repressive loops, we calculated SD-RCI with a unit of TAD (the SD-RCI calculation process can be found in Methods section) and divided the genome structure into four levels (*i*.*e*., ultra_high, high, middle, low) based on the value of d in the Gaussian distribution of SD-RCI (Figure 4B, E). To further explore the relationship between the number of RCIs and the volume of chromatin structure, we analyzed the relationship between the number of active loops or repressive loops in a topologically associated domain (TAD) and the volume of the TAD, respectively (Figure 4C, F). The results show that a structure with higher SD-RCI (*i*.*e*., high, middle level) can cover more RCIs with a smaller volume of TADs, consistent with the definition of SD-RCI.

To investigate the gene expression changes within the TADs, we calculated the gene expression value log (RPKM) for different levels of TADs from the active model. The result shows that the gene expression value is positively correlated with the SD-RCI level (Figure 4D), whereas the log (RPKM) calculated from the repress model shows an opposite change (Figure 4G). The results further proved that the repress model could detect more chromatin interactions related to inhibition of transcription, and the active model could detect more chromatin interactions related to the promotion of transcription. To explore the reason, we developed a MINE-viewer tool to do 3D structure visualization with the gene expression for the four levels of TADs using the Pastis-PM2 (Varoquaux et al., 2014) algorithm (Figure 4H, I). The results show that the spatial density of TAD in the high level is higher than in middle and low levels. Then, we visualized the loops, CTCF, H3K27ac, H3K27me3 and H3K9me3 histone ChIP-seq tracks in the active and repress high level TADs. We found that the active high level TAD regions are enriched with CTCF and H3K27ac, the repressive high level TAD regions are enriched with H3K9me3, which represses the transcriptional activity of genes (Figure 4J). The visualization of other three SD-RCI levels in active or repressive regions can be seen from Figure S16, S17, S18, S19. The above results show that RCIs called from active model or repressive model are enriched with active-related TFs or repressive-related TFs.

Based on the active or repressive loops identified by active model or repressive model, the genome regions can be spatially divided into active and repressive hubs, with active hubs in regions enriched with active TFs (*e*.*g*., CTCF, POLR2A, and SMC3) and repressive hubs in regions enriched with transcriptional repressors (*e*.*g*., EZH2). Using SD-RCI values, active and repressive hubs can be further defined into developed hubs, which have high SD-RCI (middle, high and ultra-high level), or developing hubs, which have lower SD-RCI (SD-RCI level is low) (Figure 4K). Hubs form in chromatin regions through active or repressive loops with TFs. Active or repressive hubs form where corresponding regulatory elements anchor to ensure the transcription or repression of genes required or not, respectively, for organismal function. When the proportion of RCIs in the chromatin space increases (*i*.*e*., forms a developed hub), then there is a higher frequency of anchoring by the corresponding regulatory element at a gene requiring its regulation. When the proportion of regulatory elements in the chromatin space is low (*i*.*e*., in developing hubs), the frequency of the required regulatory interaction is lower. So, genes in these developing hubs are upregulated more slowly, or require additional recruitment factors, while repression from an active state is also slower. This conclusion is consistent with the results that the average gene expression is higher in a higher SD-RCI level in the active model, and the average gene expression is lower in a higher SD-RCI level in the repressive model, as shown in Figure 4D, G.

In conclusion, the MINE-density and MINE-Viewer tools provide us with a view of the 3D visualization of spatial density of regulatory chromatin interactions, and allow us to explore the active and repressive genome by calculating the SD-RCI of active and repressive interactions, respectively.

### Spatial regulatory chromatin interaction density reflects changes in chromosome structure

MINE was next applied to data obtained from a liquid-liquid phase separation (LLPS) experiment (Ulianov et al., 2021) reported in 2021 to verify whether MINE could identify the effects of 1,6-hexanediol (1,6-HD)-mediated LLPS disruption on RCIs (Figure 5, Figure 6). For this analysis, Hi-C and epigenomic data obtained from the HeLa cell line treated (Hex group) or not (control group) with 1,6-hexanediol were downloaded (Table S1) and processed according to the MINE workflow described above.

**Figure 5.**
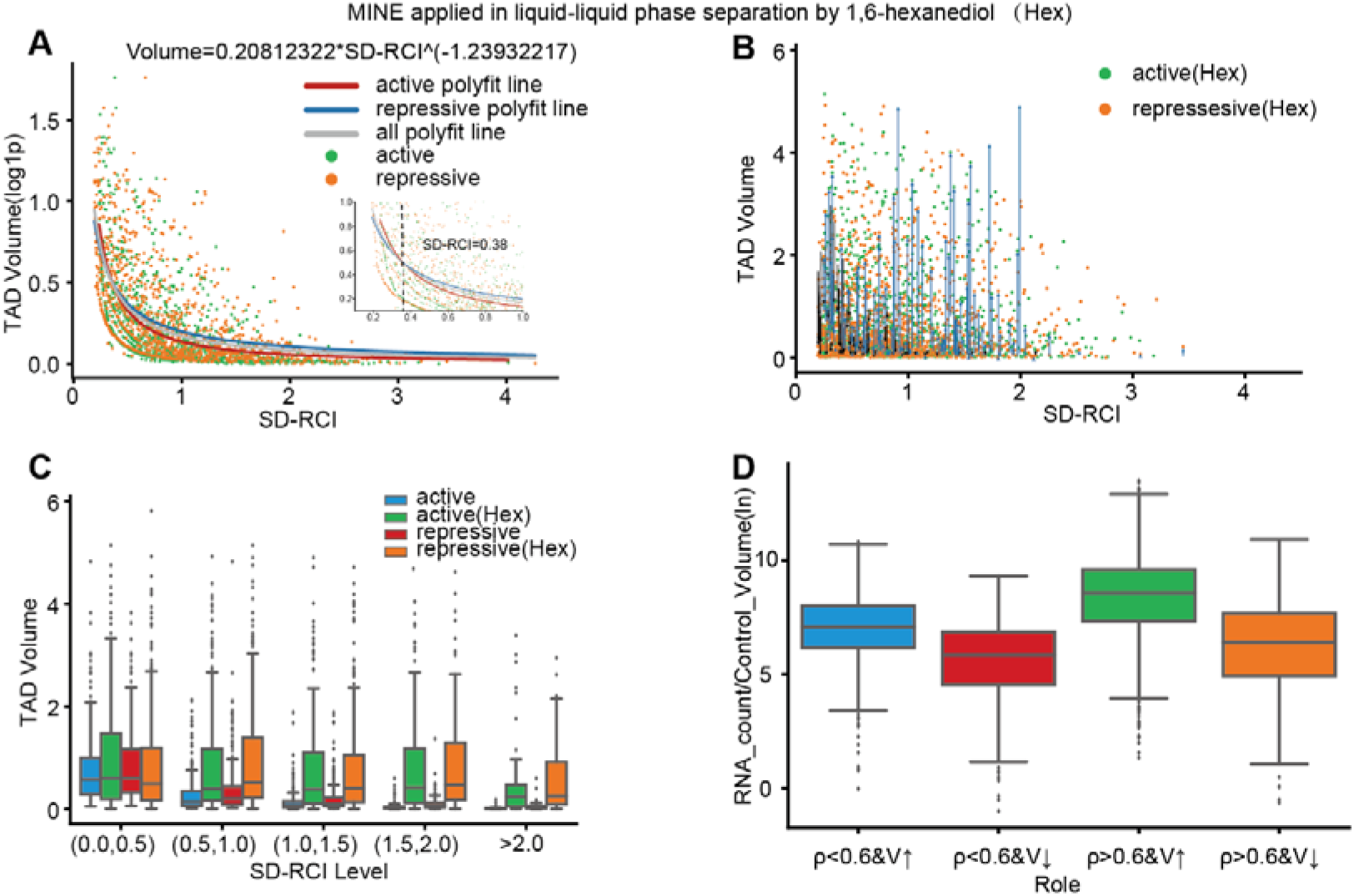
SD-RCI is correlated with chromosome structure before and after the effects of 1,6-hexanediol (1,6HD)-mediated LLPS disruption. (A–B) The distribution of the volume of TADs changes with the SD-RCI value before and after liquid-liquid phase separation. (C) The box plot of the volume of TADs and the range of SD-RCI. (D) The box plot of the gene counts ratio calculated from RNA-seq data (control group) and the four types, where p represents the value of SD-RCI, and V represents the volume of TADs.

**Figure 6.**
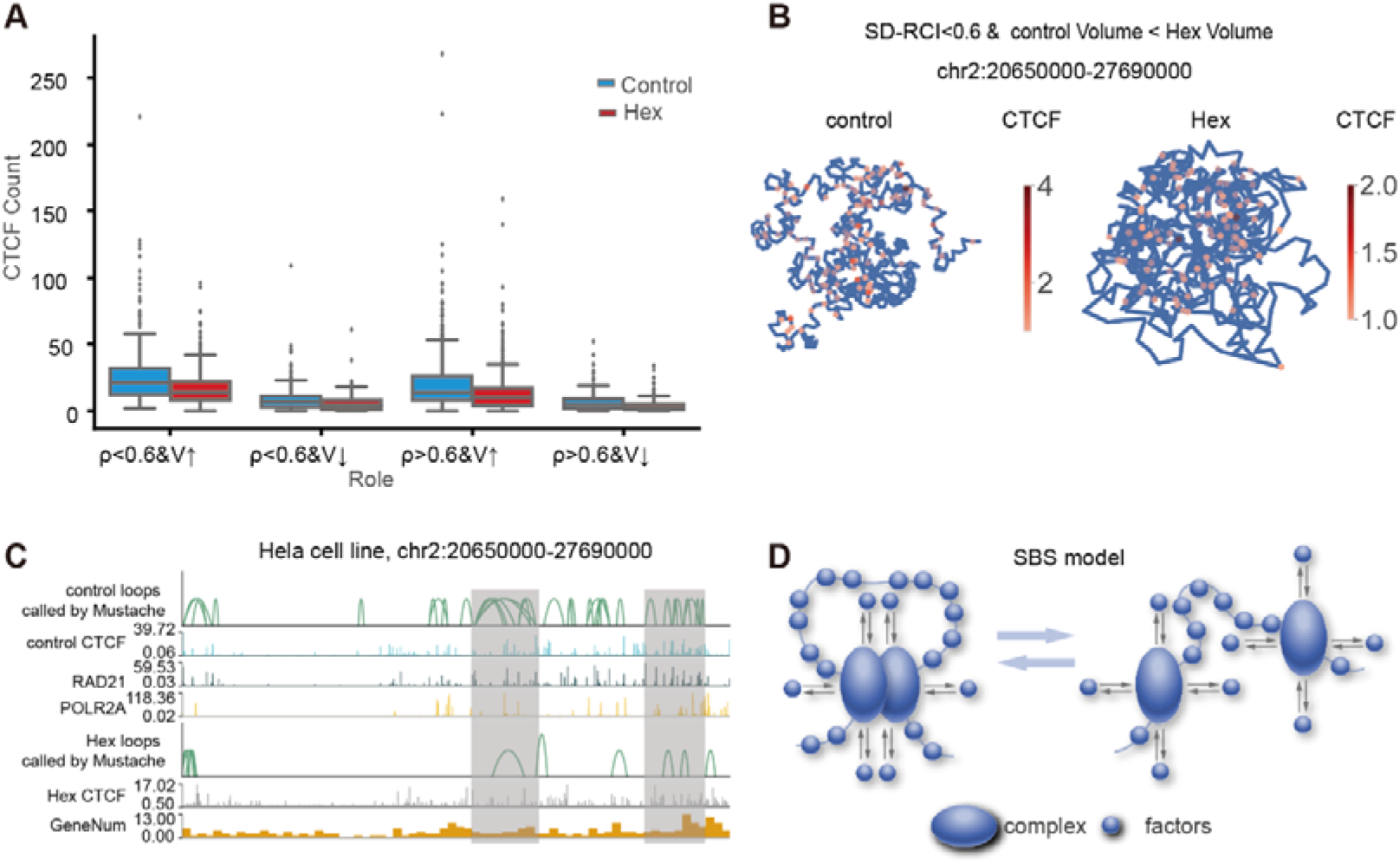
Exploration of mechanisms to explain the reason for changes in TAD volume. (A) The box plot of the CTCF counts calculated from control group and Hex group. (B) The 3D structure marked with CTCF anchor intensity of TAD typed with “SRED<0.6 & control Volume << Hex Volume”. (C) The visualization of loops and tracks from control group and Hex group. (D) the mechanism of chromatin interaction formation and unwinding.

First, we examined the relationship between these TAD volumes and SD-RCI values calculated from the active and repressive models before and after phase separation, and fitted the corresponding power function curve. We found that the active volume was larger than the repressive volume under the same SD-RCI condition when the SD-RCI of the HeLa cell line was <0.38, whereas the active volume was smaller than the repressive volume when the SD-RCI was >0.38 (Figure 5A). By comparing the volume of TADs with the active and repressive states under the same SD-RCI condition, we could determine that, for the same SD-RCI, if the volume of active TADs is larger than repressive TADs, then the number of active loops is less than the repressive loops. As we know, active regulatory interactions promote higher gene expression and repressive regulatory interactions repress gene expression. Therefore, when SD-RCI is lower than the intersection value (*i*.*e*., 0.38), the active or repressive TADs both tend to have low gene expression. When SD-RCI is higher than the intersection value (*i*.*e*., 0.38), the active or repressive TADs both tend to have high gene expression.

Enhanced Hi-C data revealed that TAD volume was larger after 1,6-HD treatment than before treatment under the same SD-RCI condition (Figure 5B, C). TADs were then categorized into four types (①SD-RCI < 0.6 and control volume < Hex volume, ②SD-RCI >= 0.6 and control volume > Hex volume, ③SD-RCI < 0.6 and control volume > Hex volume, ④SD-RCI >= 0.6 and control volume < Hex volume, where 0.6 was the SD-RCI inflection point when count became positive calculated from the count - SD-RCI curve as shown in Figure S20) according to the change in TAD volume (whether increased or decreased) and the size of SD-RCI before and after drug treatment.

To further determine whether changes in volume from pre- to post-phase separation were related to the intensity of gene transcription, we calculated gene counts ratio (*i*.*e*., *count* / | *V*_*before*_ − *V*_*after*_ |, where count is gene count, *V*_*before*_ is the volume before LLPS, *V*_*after*_ is the volume after LLPS) for the four types of changes in TAD volume. The results showed that TADs with high gene counts ratio were more likely to increase in volume, while TADs with low gene counts would likely decrease in volume (Figure 5D). Previous studies (Stadhouders et al., 2019) (Barutcu et al., 2015; Ryba et al., 2010) have established that regions with a high density of genes are mostly in open chromatin regions (A compartment), while regions with low gene density are generally in closed chromatin regions (B compartment). We therefore concluded that in the A compartment, the volume of TADs was more likely to increase following 1,6-HD treatment, while the volume of TADs was likely to decrease in the B compartment after 1,6-HD treatment. This conclusion is consistent with the established findings (Liu et al., 2021; Ulianov et al., 2021).

To explore the reason of the volume change after 1,6-HD treatment, we calculated the CTCF count distribution of the four types of TADs and found TADs with high CTCF counts were more likely to increase in volume (Figure 6A). The four types of TADs were then visualized as three-dimensional structures marked with CTCF anchor intensity before and after 1,6-HD treatment (CTCF intensity is shown in Figure 6B and Figure S21 in Supplementary Files 2). Set the type of “SRED<0.6 & control Volume << Hex Volume” as an example, the visualization of 3D structure and loops showed the control group obtained more loops than the Hex group (Figure 6C). The visualization of other three type can be found in Figure S22, S23, S24 (Supplementary Files 2). This means that the 1,6-HD treatment breaks or forms some loops, which makes the volume of TAD increase or decrease. In the Discussion section, we will use the “Strings and Binders Switch” (SBS) (Figure 6D) (Barbieri et al., 2012; Fiorillo et al., 2020) model to describe the reason of increase in chromosome volume.

## DISCUSSION

In consideration of these regulatory chromatin interactions identified through MINE-Loop, we propose that the spatial density of regulatory chromatin interactions (SD-RCI) can serve as a metric for quantitative exploration of the relationship between active or repressive chromatin interactions and the gene transcriptional status for a given TAD region. In this work, we defined four levels of SD-RCI (ultra_high, high, middle, low) to assess the relationship between SD-RCI and gene transcription. By comparing the expression strength of genes at different SD-RCI levels in the HepG2 cell line, we found that a higher SD-RCI in the active model is more conducive for gene transcription, and conversely in repressive model, higher SD-RCI is associated with greater transcriptional inhibition.

In analyses investigating the relationship between SD-RCI and chromosomal structure, SD-RCI values were used to compare HeLa cell volumes before and after liquid-liquid phase separation (LLPS) (*i*.*e*., the HeLa cell line treated (Hex group) or not with 1,6-hexanediol (control group)). We found that the nucleus volume increased following LLPS. We proposed the “Strings and Binders Switch” (SBS) (Barbieri et al., 2012; Fiorillo et al., 2020) model can explain this change in volume (Figure 6B) through the formation of loops and domains resulting from chromatin contacts between distant loci mediated by molecular factors, such as transcription factors (TFs). Before treating the HeLa cell with 1,6-hexanediol (Control group), the density of CTCF factor in 3D structure is higher than in Hex group (Figure 6A, C), a stable chromatin loop is formed through entropic force (Cook and Marenduzzo, 2018) exerted by other small molecular factors in the nucleus localized in complexes attached to the chromatin. After treating the HeLa cell with 1,6-hexanediol (Hex), the density of CTCF factor in 3D structure is lower than in Control group (Figure 6A, C), the complex is no longer attached to the same chromatin location, and the small molecular factors in the nucleus exert entropic force on each complex individually, resulting in disruption of the previously stable chromatin loops, loosening the chromatin, and resulting in increased volume. Previous studies have shown that TAD structures and the A/B compartments are primarily formed by chromatin loops extrusion, (Wit, 2019) ultimately resulting in a higher nucleus volume after treatment with 1,6-hexanediol.

In summary, we established a framework by integrating multiple omics datasets (*i*.*e*., ATAC-seq and ChIP-seq) to reduce noise and increase the proportion of detectable regulatory chromatin interactions. Applying the MINE pipeline to explore the relationship between the spatial density of regulatory chromatin interactions (SD-RCI) and gene transcriptional status led to the discovery of four levels of spatial density in chromatin interactions that reflect the relationship between SD-RCI and gene regulation. We then applied MINE to data obtained from a liquid-liquid phase separation experiment (i.e., treating the HeLa cell with 1,6-hexanediol), which showed that the 3D conformation of active and repressive models are consistent with the results that the 1,6-HD treatment caused the enlargement of nucleosome clutches and their more uniform distribution in the nuclear space (Ulianov et al., 2021). Finally, the mechanism underlying structural changes in TADs before and after LLPS was explained by the Strings and Binders Switch (SBS) model (Barbieri et al., 2012; Fiorillo et al., 2020).

## STAR*METHODS

Detailed methods are provided in “Supplementary Files 1”.

## Data availability

The analysis code is available in the GitHub repository (https://github.com/MICL-biolab/MINE).

## SUPPLEMENTAL INFORMATION

Supplementary data are available online at bioRxiv.

## Acknowledgements

This work was funded by grants from the National Key R&D Program of China (2018YFB0704301, 2018YFB0704304, 2018YFA0801402), the Scientific and Technological Innovation Foundation of Shunde Graduate School, USTB (BK20BF009), the National Natural Science Foundation of China (31871343,61971031) and the CAMS Innovation Fund for Medical Sciences (2020-RC310-009). Funding for open access charge: Department of Computer Science and Technology, Advanced Innovation Center for Materials Genome Engineering, University of Science and Technology Beijing. The authors wish to thank Chen Fengling in Tsinghua university for her suggestions about data analysis, thank Xu miaomiao in University of Science and Technology Beijing for her assistance on the color and layout of figures in the manuscript.

## AUTHOR CONTRIBUTIONS

All authors contribute equally.

## DECLARATION OF INTERESTS

The authors declare that they have no competing interests.

## Notes

https://github.com/MICL-biolab/MINE

### Competing Interest Statement

The authors have declared no competing interest.

