## Supplementary Files 1 for "MINE is a method for detecting spatial density of regulatory chromatin interactions based on a MultI-modal NEtwork"

^1^School of Computer and Communication Engineering, Beijing Key Laboratory of Knowledge Engineering for Materials Science, Beijing Advanced Innovation Center for Materials Genome Engineering, University of Science and Technology Beijing, Beijing100083, China. ^2^The State Key Laboratory of Medical Molecular Biology, Department of Biochemistry and Molecular Biology, Institute of Basic Medical Sciences, School of Basic Medicine, Chinese Academy of Medical Sciences, Peking Union Medical College, Beijing 100005, China. ^3^ Shunde Graduate School, University of Science and Technology Beijing, Foshan 528399, China. ^4^ School of mechanical engineering, University of Science and Technology Beijing, Beijing100083, China.

Haiyan Gong and Minghong Li contributed equally to this work.

**METHODS**

### Datasets

The description of Hi-C data. Hi-C data for six human cell lines (GM12878, IMR90, K562, H1-hESC, HepG2, and Hela) are downloaded from https://data.4dnucleome.org/ (Dekker et al., 2017) under accession number 4DNFI1UEG1HD, 4DNFIH7TH4MF, 4DNFITUOMFUQ (Rao et al., 2014), 4DNFI2TK7L2F (Krietenstein et al., 2020), 4DNFICSTCJQZ, and 4DNESCMX7L, respectively.

The description of epigenome data that are used for training. For the active model, ATAC-seq data and ChIP-seq data of H3K27ac and H3K4me3 mainly target transcription-related factors are selected, for the repressive model, ChIP-seq data of H3K9me3 and H3K27me3 that are associated with gene repression are selected.

To verify that the MINE-Loop method can help identify more regulatory chromatin interactions, the CTCF, RAD21, SMC3, POLR2A ChIP-seq data are selected to verify the gene transcription-related interaction, and POLR2A, EZH2 ChIP-seq data are selected to verify the gene repression-related interactions. The download links of all data can be obtained from Table S1 in the Supplementary Files2.

To verify whether the loops called from the MINE_enhanced_hic can overlap the loops called from ChIA-PET data, CTCF and POLR2A ChIA-PET data in HepG2 cell line were obtained from ENCODE (Consortium, 2004) under accession number ENCSR411IVB and ENCSR857MYZ, respectively.

### Down sampling Hi-C data

Downsampling Hi-C matrix were simulated by downsampling the VC-normalized Hi-C matrix obtain from .hic format file using juicerbox (Robinson et al., 2018). The downsampling strategy is described as Figure S14:
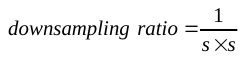
 is defined, then, values of the
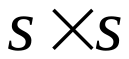
 window were all set as the average value of the
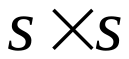
window.

### Complete downsampling Hi-C matrices

To obtain more Hi-C matrix features, we first synthesize these downsampling Hi-C matrices by point-to-point addition, and then complete the Hi-C matrix after addition through the FAN method (Achanta et al., 2017), a matrix completion algorithm suited for the sparse input matrix whose 99% pixels are randomly missing. For the FAN method, we first define the row or column of the matrix after superimposing as f, define the signal as g containing ones and zeros with similar length of f, as described in Eq. (1). Then, f and g are convolved with the same kernel h. Finally, we can obtain the completed matrix by performing an element-wise division of the two convolved signals, as Eq. (2) shows.

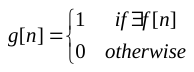
（1）

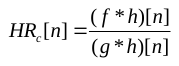
（2）

Where n represents the position of the signal, g[n], f[n], and
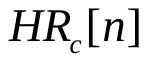
 represent the signal g, f, and the finally obtained complete Hi-C data at position n, respectively.

### Preproces epigenome data (Generation of Correlation matrix)

Same as Hi-C matrix, we divided the chromosome into n fragments with a length of len (in this paper, len=1000 base), and calculated the signal p-value
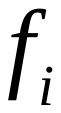
 of the *ith* fragment using ChIP-seq or ATAC-seq data as the following formula to get a
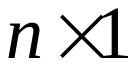
 matrix.

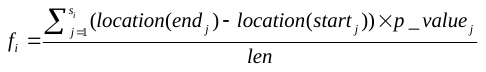
（3）

Where
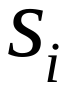
 is the number of fragments divided by ChIP-seq data within the fragment i,
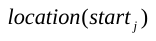
 and
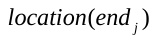
 is the start and end location of the jth fragment in ChIP-seq data,
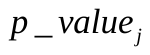
 is the signal p-value of the jth fragment in ChIP-seq data.

Then we combine multiple matrixes in columns to obtain a feature matrix
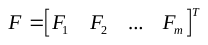
 and normalize it to be 0~1with
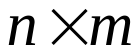
 size, where m is the sample number of the epigenomics data. We define the i-th row of feature matrix F as
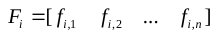
, the average of matrix F as
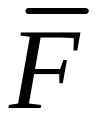
,
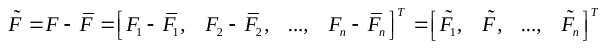
, the transpose of
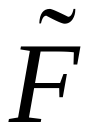
as
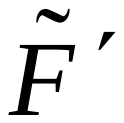
,
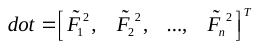
, then the Pearson correlation coefficient between pairwise of F can be calculated as
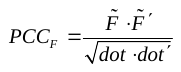
, where
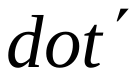
is the transpose of
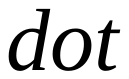
.

### Matrix Normalization

For the data used in the training, **the completed matrix** obtained from section “Complete downsampling Hi-C matrices”, **Pearson correlation coefficient matrix** obtained from section “Preproces epigenome data (Generation of Correlation matrix)” and **VC-normalized Hi-C matrix** obtain from .hic format file using juicerbox (Robinson et al., 2018) all need to be normalized as Eq. (4-7) shows. We define
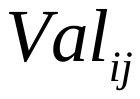
 as the value of row i and column j, num pairs as the number of pairs whose value is not zero, nums as the 1/1000 of the sum of num pairs as Eq. (1),
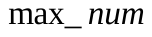
as the minimum value of the largest nums value in the matrix. We first set the values greater than
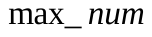
 to be
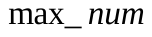
 for these matrices, then do normalization as Eq. (6, 7) to limit the value to be between 0 and 255.

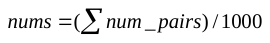
（4）

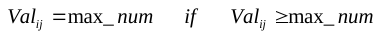
（5）

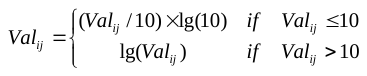
（6）

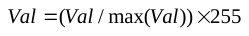
（7）

#### Generation of Masked_hic

In order to ensure that the MINE-Loop method can obtain more regulatory chromatin interactions, the VC-normalized Hi-C matrix after normalization as Eq. (4-7) was masked with the Candidate Cis-Regulatory Elements or other ChIP-seq data to be the target high-resolution Hi-C used in training **(Masked_hic).** The mask operation obeys the following principle: for ChIP-seq data, only these interaction values with peaks in the ChIP-seq data at both positions will be retained; for the cis-regulatory element, we first set values of all locations in the cis-regulatory element file to be 1, then retain the interaction values of Hi-C matrix when values of both ends in the cis-regulatory element file are 1.

With the above operations, we can obtain all data for training: completed Hi-C (**Completed_hic**), Pearson correlation coefficient matrix (
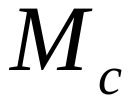
) and the target high-resolution Hi-C (**Masked_hic**).

In order to compare with the raw high-resolution Hi-C matrix at the same scale, the raw high-resolution Hi-C matrix was also normalized to be 0-255 as Eq. (4, 5, 7). In the remaining manuscript, we name **the Hi-C used for comparison with the enhanced Hi-C as Raw_hic**.

### Matrix-based sample division

Divide the Masked_hic, Completed_hic and
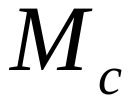
 into
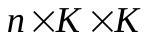
sub-matrices, where n represents the number of sub-matrices, K is the dimension of sub-matrix (in this paper, we define K=400), each sub-matrix is treated as a sample. The reason for choosing a
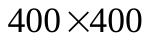
size is that according to the previous data enhancement algorithm HiCPlus (Yan et al., 2018), the interaction within the genomic distance of
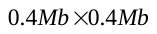
can retain more local information.

As shown in Figure S15, since the Hi-C matrix is a symmetric matrix, only the upper right part of the diagonal is reserved. Along the upper right corner of the Hi-C matrix, sub-matrices of
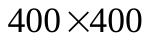
are taken as samples from top to bottom in a step of 400 kb. Each small square in the figure represents a
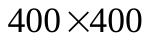
sub-matrix, and 5 sub-matrices are taken (that is, the two interaction sites are within the 2Mb genome range, because the average size of TADs is within the 1Mb genome distance range. Outside of TADs, few significant interactions are existing).

### Structure of the MINE-loop network

The implementation of the network is shown in Figure S1. The network structure is divided into three type layers: MINE_Conv, maxPool2D, and ConvTranspose_2D as Eq. (8-11) show. The first type (*i.e.*, MINE_Conv in Figure S1b, containing Conv2d, BatchNorm2d, ReLU, Conv2d, BatchNorm2d, and ReLU, is proposed to extract and present the Hi-C pattern. The second (*i.e.*, MaxPool2d in Figure S1a), is designed to perform dimensionality reduction operations. The third (*i.e.*, ConvTranspose2d in Figure S1a), is designed to perform an upsampling operation. As Figure S1a shows, we divided the network’s input into two parts: the completed Hi-C data
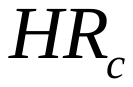
 and the correlation matrix of the epigenomics data
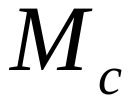
. For
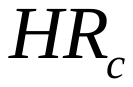
, we use MEMR_Conv Module following with a max pool layer to do downsampling, and ConvTranspose 2D layer to do upsampling times to get a result (c5). For
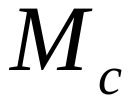
, we use MINE_Conv Module three times to get a result (e3) and merge c5 and e3, and put them into the network (MINE_Conv Module two times) to get the final enhanced Hi-C matrix.

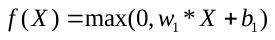
（8）

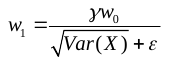
（9）

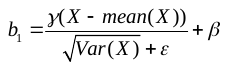
（10）

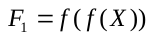
（11）

where
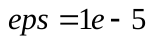
,
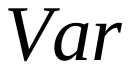
represents the variance,
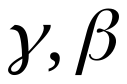
 represent the learned coefficient matrix gamma and beta,
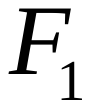
 represents the output of MINE_Conv.

### Model training and testing for MINE-loop

Data (**Completed_hic, correlation matrix and Masked_hic**) of chromosome 1-17 in GM12878 cell line are used for training the model in this paper, data of chromosome 18-22 in GM12878 are used for testing, and the ChIP-seq data of transcription factors (RAD21, SMC3, POLR2A) from GM12878, IMR90, K562, H1-hESC and HepG2 cell lines are used for verification. The download links of all data can be obtained from Table S1 of the Supplementary Files2.

**MINE_enhanced_hic matrix and Masked_hic matrix** are used as the input of L1 Loss (Eq.12) and Perceptual Loss (Johnson et al., 2016) (Eq.13) for comparison and scoring. Since MINE_enhanced_hic may be very sparse in some subgraphs, this will affect the judgment of the Perceptual Loss on the result and indirectly affect the training result. Therefore, we remove these training data, if the number of value points of the attention interaction sub-matrix is less than 10% of the sub-matrix scale.

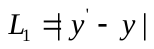
（12）

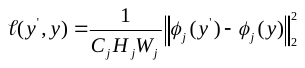
（13）

Where
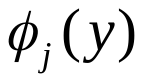
 is a feature map of shape
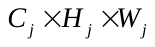
, the loss is a loss using the squared, normalized Euclidean distance between features.

As Table 1 shows, MINE-loop can be divided into active model and repress model with different epigenome data as the input of model training and Masked_hic. For the active model, ATAC-seq, H3K27ac, H3K4me3 ChIP-seq are chose to be epigenome data to train model; For the repressive model, H3K27me3, H3k9me3 ChIP-seq are chose to be epigenome data to train model. For the training of active model, we chose the cis-regulatory element file as the data source of the attention matrix for training, since there are many attention interactions and the model can be fully trained, we choose the best result when the loss score of the test set is the lowest. For the training of repressive model or when the number of attention points in the matrix is too small, we believe that the model is prone to functional overfitting due to too few comparison points. Therefore, we use the iterative version model of the test dataset with the lowest validation loss score as the final trained model.

Table 1. Data used to train the active model or repressive model.

| Model | epigenome data (input) | Data used to generate masked_hic |
| --- | --- | --- |
| Active | ATAC-seq, H3K27ac, H3K4me3 ChIP-seq | cis-regulatory element file |
| Active | ATAC-seq, H3K4me3 ChIP-seq | cis-regulatory element file |
| Active | ATAC-seq, H3K27ac, ChIP-seq | cis-regulatory element file |
| repressive | H3K27me3, H3K9me3 ChIP-seq | H3K27me3, H3k9me3 ChIP-seq |

### Evaluation methods

In this paper, we mainly verify whether the MINE_enhanced_hic matrix can be used to detect more regulatory chromatin interactions than from the Raw_hic matri from the following aspects:

**Verify Biologically:**

1) for the active model aimed at enhancing the interactions related to the promotion of transcription, we choose to explore the overlap number of loops identified by MINE_enhanced_hic and Raw_hic within different genomic distance range (2-100kb, 2-300kb and 2-500kb genomic distance), the number of CTCF, RAD2, SMC, POLR2A TFs, and transcription start site (TSS) anchoring around loops called from MINE_enhanced_hic and Raw_hic, to verify the MINE-Loop method can help to identify more loops related to the promotion of transcription.

2) Verify the influence of different types of ChIP-seq data combinations on the effect of MINE-Loop: study the influence of the model obtained by combining different epigenome data used for MINE-Loop training input on the data enhancement effect.

3) Verify whether MINE_enhanced_hic can enrich more functional genes: KEGG enrichment analysis was performed on immune-activated cells (human B lymphocyte line GM12878) and human liver cancer cell line (HepG2) to verify whether MINE_enhanced_hic can enrich more functional genes.

**Verify the general applicability of MINE-Loop model:**

1) the ability to do prediction in other cell lines data by using the active model trained by the GM12878 cell line.

2) the ability to do prediction with different combinations of histone target ChIP-seq data as the prediction input.

#### Call loops using FitHiC2 and mustache

Loops from Hi-C data are called by FitHiC2(Kaul et al., 2020) with parameters as the following: resolution = 1kb, distLowThres=2kb, and distUpThres=100kb, 300kb or 500kb, p-value<0.015, q-value (FDR obtained by applying Benjamini-Hochberg correction to the p-values) <0.015.

For the mustache (Ardakany et al., 2020) tool, loops from raw high-resolution Hi-C data were called from the .hic format file with default parameters. loops from MINE_enhanced_hic matrix were called from text format with parameters as the following: resolution = 1kb, pt (P-Value threshold) =0.5.

#### Overlap of Raw_hic and MINE_enhanced_hic

Both ends of loops called separately from Raw_hic and MINE_enhanced_hic are anchored within 2kb genomic distance are defined as overlapping.

#### Call loops using ChIA-PET2

Loops from ChIA-PET data are called by ChIA-PET2 tool (Li et al., 2016) with parameters and limitations as the following: -m 1 -A ACGCGATATCTTATC -B AGTCAGATAAGATAT.

#### Analysis of loops anchoring transcription factor

Calculate counts of CTCF, RAD21, SMC3, POLR2A, EZH2 CTCF, RAD21, SMC3 peak around loops (distance to the loop anchor point: -40kb~+40kb) called from Raw_hic and MINE_enhanced_hic. curves of factors peak count with distance to loop anchor point were plotted using python library matplotlib (Hunter, 2007).

#### gene ontology and pathway enrichment analysis in differential loop anchors

Gene KEGG pathway enrichment analysis were performed using R package org.Hs.eg.db (Carlson et al., 2019)(3.14.0), clusterProfiler (Wu et al., 2021) (4.0), dplyr (Yarberry, 2021) (1.0.7) and ggplot2 (Wickham et al., 2012) (3.3.5).

### Spatial regulatory element density

The calculation steps of SD-RCI are as follows: (1) the Pastis-PM2 (Varoquaux et al., 2014) algorithm was used to get the 3D coordinates of these bins for the TADs; (2) calculate the volume of
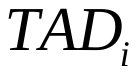
 (
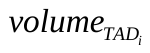
) constructed by the 3D coordinates of these bins as the raw volume of
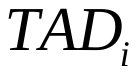
; (3) calculate the raw SD-RCI of
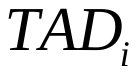
:
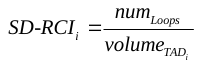
, where
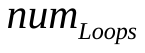
is the total number of loops in each TAD region. (4) the number of total loops may increase or decrease as sequencing depth or loop-calling algorithms changes. Same as SDOC (Jiang et al., 2021), we used quantile normalization to normalize the raw SD-RCI value to Gaussian distribution (mean = 0, standard deviation = 1). (5) the SD-RCI of the ith TAD in a chromosome predicted by HiCDB (Chen et al., 2018) can be formulated as follows.

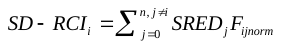
（13）

Where n is the number of TADs predicted by HiCDB (Chen et al., 2018),
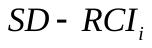
 is the ith TAD on a same chromosome,
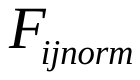
 is the TAD-TAD normalized contact frequency between
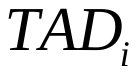
 and
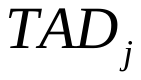
 can be formulated as follows:

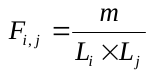
（14）

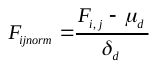
（15）

Where
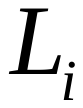
,
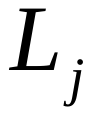
 represent the length of
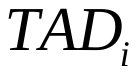
,
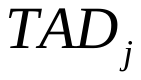
 in a chromosome, m is the sum of contact frequency between
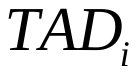
 and
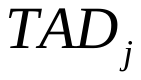
,
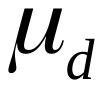
 is the loess regressed pairwise contact frequency,
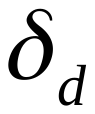
 is the loess regressed standard deviation calculated by the lowess function of statsmodels Python library. We let
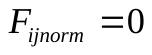
if genomic distance
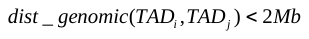
 or
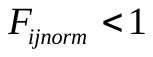
.

#### Expression changes associated with level changes in HepG2 cell line

Genes were classified based on the levels (ultra_high, high, middle, low) divided by SD-RCI in HepG2 cell line. The counts file of RNA-seq data in the HepG2 cell line was obtained from GEO under accession number GSE117815, RPKM value was calculated as the following formula.

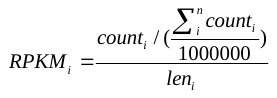
（14）

Where
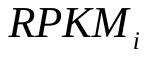
 is the ith gene’s RPKM value,
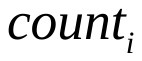
 is the ith gene’s count number,
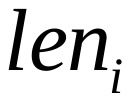
 is the length of the ith gene.

### Important definitions in MINE work

**1) Hi-C:** High-through chromosome conformation capture.

**2) RCI:** regulatory chromatin interaction.

**3) Raw_hic:** the raw high-resolution Hi-C matrix calculated from deeply sequenced Hi-C data using VC-normalized method used to do comparative analysis.

**4) Masked_hic:** the 1 kb high-resolution Hi-C matrix from raw high-resolution Hi-C masked by location of interest like candidate Cis-Regulatory Elements.

**5) MINE_enhanced_hic:** the 1 kb resolution Hi-C matrix predicted by MINE method.

**6) Completed_hic:** Hi-C matrix completed from many downsampling Hi-C matrix with different downsampling ratio.

**7) active model:** The MINE-Loop model trained using ATAC-seq data and ChIP-seq data for different targeted factors (e.g., H3K4me3 and H3K27ac) specifically involved in DNA transcription.

**8) repressive model:** The MINE-Loop model trained using ChIP-seq data target with suppression-related epigenomic marks (i.e., H3K27me3, H3k9me3).

**9) active loops:** loops identified from the active model.

**10) repressive loops:** loops identified from the repressive model.

**11) SD-RCI**: the spatial density of regulatory chromatin interactions, i.e., the ratio of the total number of active or repressive interactions in a TAD to the entire 3D space taken up by the physical structure of the TAD.

**12) Four levels of TADs’ SD-RCI value**: ultra-high, high, middle and low based on the value of δ in the Gaussian distribution of SD-RCI.

**13) The TADs were categorized into four types** (①SD-RCI < 0.6 and control volume < Hex volume, ②SD-RCI >= 0.6 and control volume > Hex volume, ③SD-RCI < 0.6 and control volume > Hex volume, ④SD-RCI >= 0.6 and control volume < Hex volume, where 0.6 was the SD-RCI inflection point when count became positive calculated from the gene count - SD-RCI curve as shown in Figure S18) according to the change in TAD volume (whether increased or decreased) and the size of SD-RCI before and after drug treatment.

**14) active hubs:** TAD regions that are enriched with active factors.

**15) developed active hubs:** TAD regions that are enriched with active factors, and the SD-RCI level is middle, high and ultra-high.

**16) active hubs:** TAD regions that are enriched with active factors.

**17) developed active hubs:** TAD regions that are enriched with active factors, and the SD-RCI level is middle, high and ultra-high.

**18) developing active hubs:** TAD regions that are enriched with active factors, and the SD-RCI level is low.

**19) repressive hubs:** TAD regions that are enriched with repressive factors.

**20) developed repressive hubs:** TAD regions that are enriched with repressive factors, and the SD-RCI level is middle, high and ultra-high.

**21) developing repressive hubs:** TAD regions that are enriched with repressive factors, and the SD-RCI level is low.
