## Supplementary Files 2 for "MINE is a method for detecting spatial density of regulatory chromatin interactions based on a MultI-modal NEtwork"

Table S1 the dataset used in MINE work.

| Cell line | motivation | Data type | Data link |
| --- | --- | --- | --- |
| GM12878 | Genral | Hi-C | <https://data.4dnucleome.org/files-processed/4DNFI1UEG1HD/> |
|  | Train and predict for Active model | ATAC-seq | <https://www.encodeproject.org/experiments/ENCSR637XSC/> |
|  |  | H3K27ac ChIP-seq | <https://www.encodeproject.org/experiments/ENCSR000AKC/> |
|  |  | H3K4me3 ChIP-seq | https://www.encodeproject.org/experiments/ENCSR057BWO/ |
|  |  | Cis-Regulatory Elements | https://www.encodeproject.org/annotations/ENCSR820WFY/ |
|  | Train and predict for repress model | H3K27me3 ChIP-seq | https://www.encodeproject.org/experiments/ENCSR000DRX/ |
|  |  | H3K9me3 ChIP-seq | https://www.encodeproject.org/experiments/ENCSR000AOX/ |
|  | validation | CTCF ChIP-seq | https://www.encodeproject.org/experiments/ENCSR000DKV/ |
|  |  | RAD21 ChIP-seq | https://www.encodeproject.org/experiments/ENCSR000BMY/ |
|  |  | SMC3 ChIP-seq | https://www.encodeproject.org/experiments/ENCSR000DZP/ |
|  |  | POLR2A ChIP-seq | https://www.encodeproject.org/experiments/ENCSR000EAD/ |
|  |  | EZH2 | https://www.encodeproject.org/experiments/ENCSR000ARD/ |
|  |  | Annotation file | ftp://ftp.ensembl.org/pub/release-104/gff3/homo_sapiens/Homo_sapiens.GRCh38.104.chr.gff3.gz |
| H1-hESC | Train and predict | Hi-C | https://data.4dnucleome.org/files-processed/4DNFI2TK7L2F/ |
|  |  | ATAC-seq | https://data.4dnucleome.org/files-processed/4DNFICPNO4M5 |
|  |  | H3K27ac ChIP-seq | https://www.encodeproject.org/experiments/ENCSR880SUY/ |
|  |  | H3K4me3 ChIP-seq | https://www.encodeproject.org/experiments/ENCSR443YAS/ |
|  |  | Cis-Regulatory Elements | https://www.encodeproject.org/annotations/ENCSR597SZL/ |
|  | validation | CTCF ChIP-seq | https://www.encodeproject.org/experiments/ENCSR000BNH/ |
|  |  | RAD21 ChIP-seq | https://www.encodeproject.org/experiments/ENCSR000BLD/ |
|  |  | POLR2A ChIP-seq | https://www.encodeproject.org/experiments/ENCSR000BHN/ |
| K562 | Train and predict | Hi-C | https://data.4dnucleome.org/files-processed/4DNFITUOMFUQ/ |
|  |  | ATAC-seq | https://www.encodeproject.org/experiments/ENCSR483RKN/ |
|  |  | H3K27ac ChIP-seq | https://www.encodeproject.org/experiments/ENCSR000AKP/ |
|  |  | H3K4me3 ChIP-seq | https://www.encodeproject.org/experiments/ENCSR000AKU/ |
|  |  | H3K9me3 ChIP-seq | https://www.encodeproject.org/experiments/ENCSR000APE/ |
|  |  | H3K27me3 ChIP-seq | https://www.encodeproject.org/experiments/ENCSR000EWB/ |
|  |  | Cis-Regulatory Elements | https://www.encodeproject.org/annotations/ENCSR301FDP/ |
|  | validation | CTCF ChIP-seq | https://www.encodeproject.org/experiments/ENCSR000DMA/ |
|  |  | RAD21 ChIP-seq | https://www.encodeproject.org/experiments/ENCSR000BKV/ |
|  |  | SMC3 ChIP-seq | https://www.encodeproject.org/experiments/ENCSR000EGW/ |
|  |  | POLR2A ChIP-seq | https://www.encodeproject.org/experiments/ENCSR388QZF/ |
|  |  | EZH2 ChIP-seq | https://www.encodeproject.org/experiments/ENCSR000AQE/ |
| IMR90 | Train and predict | Hi-C | https://data.4dnucleome.org/files-processed/4DNFIH7TH4MF/ |
|  |  | ATAC-seq | https://www.encodeproject.org/experiments/ENCSR200OML/ |
|  |  | H3K27ac ChIP-seq | https://www.encodeproject.org/experiments/ENCSR002YRE/ |
|  |  | H3K4me3 ChIP-seq | https://www.encodeproject.org/experiments/ENCSR087PFU/ |
|  |  | H3K9me3 ChIP-seq | https://www.encodeproject.org/experiments/ENCSR055ZZY/ |
|  |  | Cis-Regulatory Elements | https://www.encodeproject.org/annotations/ENCSR599FOY/ |
|  | validation | CTCF ChIP-seq | https://www.encodeproject.org/experiments/ENCSR000EFI/ |
|  |  | RAD21 ChIP-seq | https://www.encodeproject.org/experiments/ENCSR000EFJ/ |
|  |  | SMC3 ChIP-seq | https://www.encodeproject.org/experiments/ENCSR000HPG/ |
|  |  | POLR2A ChIP-seq | https://www.encodeproject.org/experiments/ENCSR000EFK/ |
| HepG2 | Train and predict | Hi-C | https://data.4dnucleome.org/experiment-set-replicates/4DNESC2DEQIJ/ |
|  |  | ATAC-seq | https://www.encodeproject.org/experiments/ ENCSR042AWH / |
|  |  | H3K27ac ChIP-seq | https://www.encodeproject.org/experiments/ENCSR000AMO/ |
|  |  | H3K4me3 ChIP-seq | https://www.encodeproject.org/experiments/ENCSR575RRX/ |
|  |  | H3K9me3 ChIP-seq | https://www.encodeproject.org/experiments/ENCSR000ATD/ |
|  |  | H3K27me3 ChIP-seq | https://www.encodeproject.org/experiments/ ENCSR000AOL/ |
|  | validation | CTCF ChIP-seq | https://www.encodeproject.org/experiments/ENCSR000AMA/ |
|  |  | RAD21 ChIP-seq | https://www.encodeproject.org/experiments/ENCSR000EEG/ |
|  |  | SMC3 ChIP-seq | https://www.encodeproject.org/experiments/ENCSR000EDW/ |
|  |  | POLR2A ChIP-seq | https://www.encodeproject.org/experiments/ENCSR000EEM/ |
|  |  | EZH2 ChIP-seq | https://www.encodeproject.org/experiments/ENCSR000ARI/ |
|  |  | CEBPB ChIP-seq | https://www.encodeproject.org/experiments/ENCSR000BQI/ |
| Hela（within different condition） | Train and predict | hic | <https://data.4dnucleome.org/experiment-set-replicates/4DNESCMX7L58/> |
|  |  |  | <https://www.ncbi.nlm.nih.gov/geo/query/acc.cgi?acc=GSE138543> |
|  |  | ATAC | <https://www.ncbi.nlm.nih.gov/geo/query/acc.cgi?acc=GSE121840> |
|  |  | H3K27ac（Lab: Bradley Bernstein, Broad） | <https://www.encodeproject.org/experiments/ENCSR000AOC/> |
|  |  | H3K4me3（Lab: Bradley Bernstein, Broad） | <https://www.encodeproject.org/experiments/ENCSR000AOF/> |
|  |  | H3K27me3（Lab: Bradley Bernstein, Broad） | <https://www.encodeproject.org/experiments/ENCSR000APB/> |
|  |  | H3K9me3 (Lab: Bradley Bernstein, Broad) | <https://www.encodeproject.org/experiments/ENCSR000AQO/> |
|  | validation | CTCF_peaks (Lab: Bradley Bernstein, Broad) | <https://www.encodeproject.org/experiments/ENCSR000AOA/> |
|  |  | CTCF_peaks (Hex5) | <https://www.ncbi.nlm.nih.gov/geo/query/acc.cgi?acc=GSE138543> |
|  |  | POLR2A_peaks | <https://www.encodeproject.org/experiments/ENCSR000BGO/> |
|  |  | RAD21_peaks | <https://www.encodeproject.org/experiments/ENCSR000EDE/> |
|  |  | SMC3_peaks | <https://www.encodeproject.org/experiments/ENCSR000ECS/> |
|  |  | EZH2_peaks | <https://www.encodeproject.org/experiments/ENCSR000ATC/> |

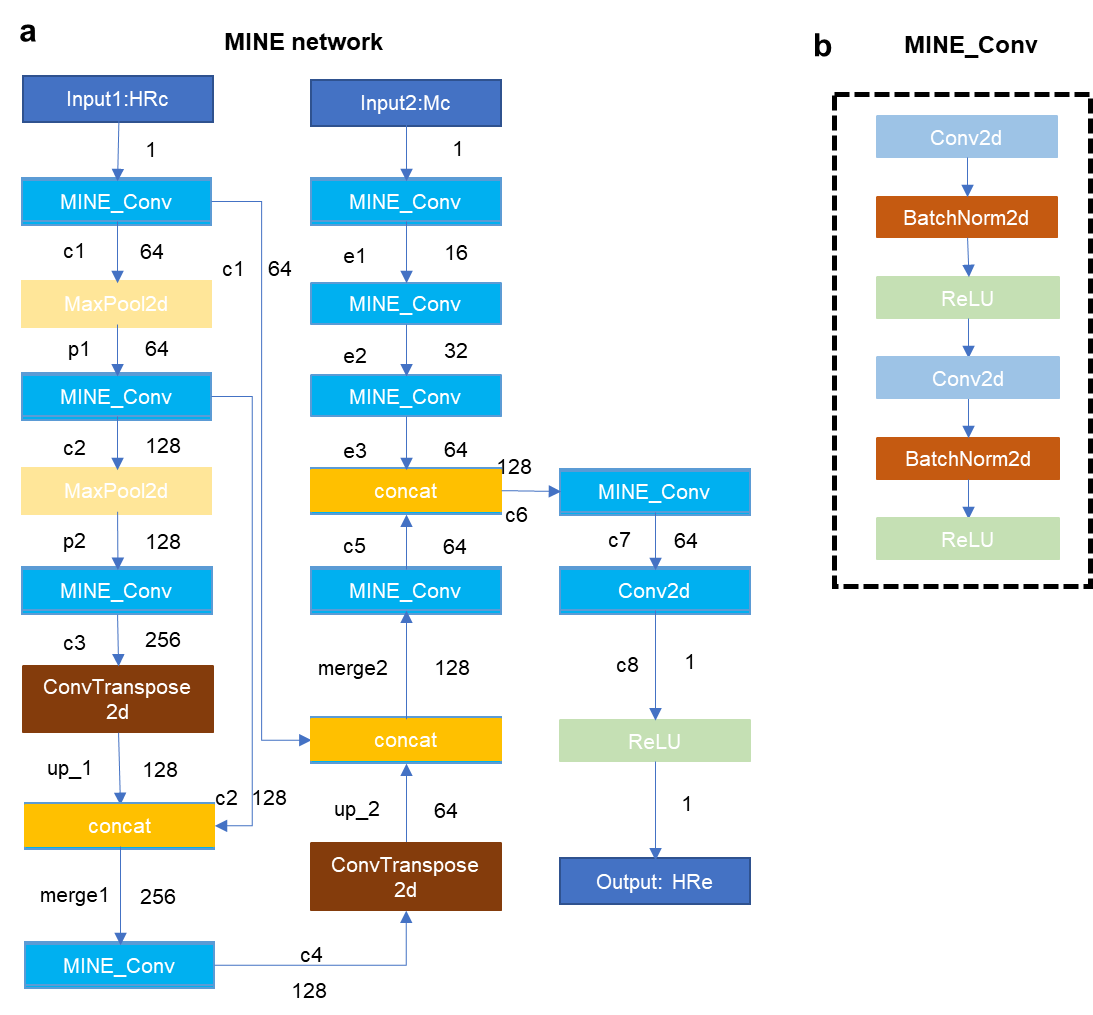

**Figure S1 –** the neural network structure of MINE-loop.

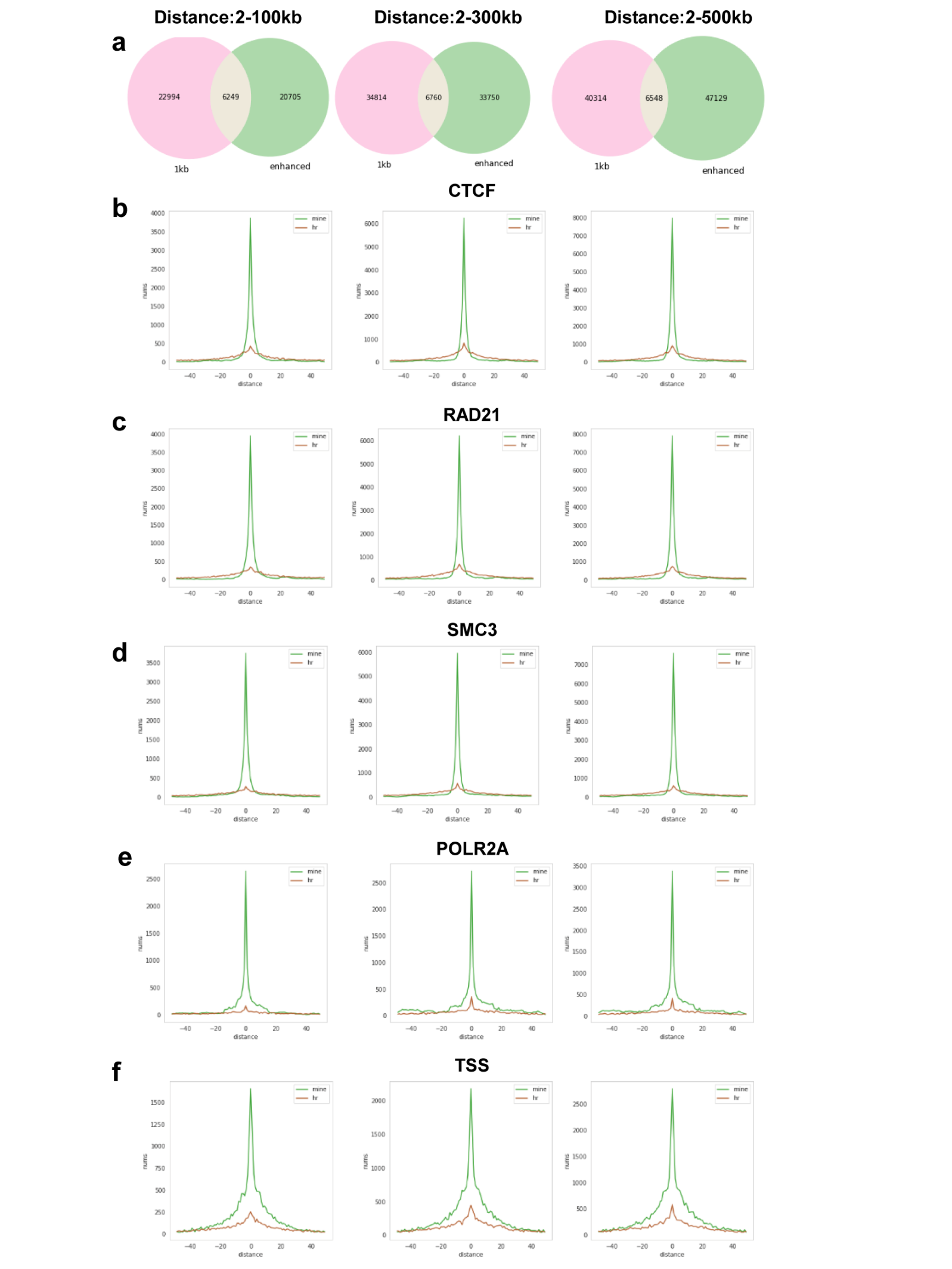

**Figure S2 –**The hit distance distribution graph for the biologically meaningful interaction using the prediction from the model trained using the completed Hi-C, ATAC-seq, H3K27ac, H3K4me3 ChIP-seq data in GM12878 cell line. The graphs in columns 1-3 represent the results within the range of 2-100, 2-300, and 2-500 genome distance. Among them, from top to the bottom are the overlapping Venn diagrams of the loops detected separately from MINE_enhanced Hi-C and the raw high-resolution Hi-C detection using the FitHiC2^1^ tool, the number of hits change curve graph of CTCF factor, RAD21, SMC3, POLR2A and TSS loci with the genomic distance, where green represents the hit from the MINE_enhanced Hi-C, and red represents the hit from the original high-resolution Hi-C.

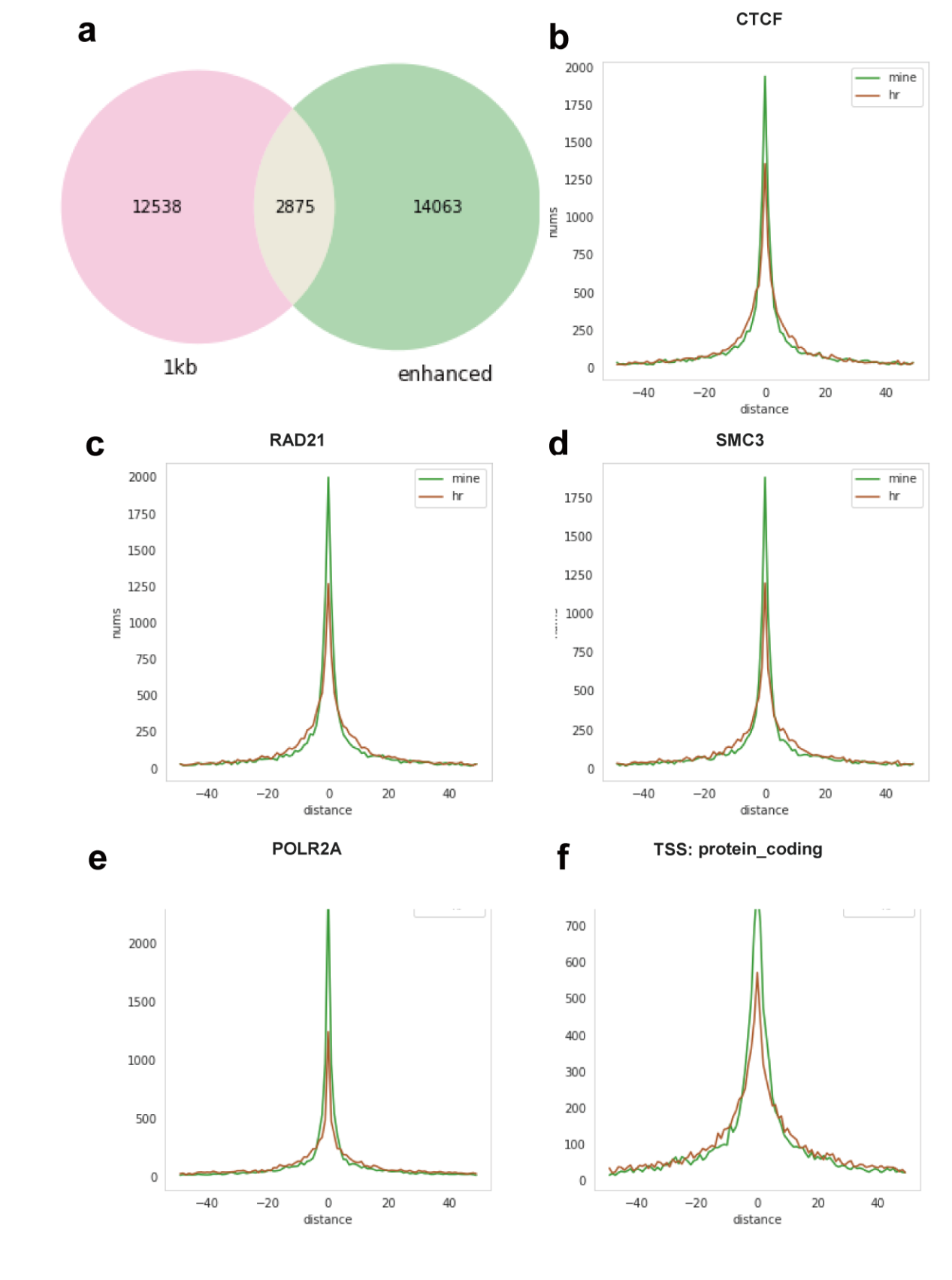

**Figure S3 –** the active model trained by GM12878 cell line, we using mustache^2^ tool to call loops, as same with FitHiC2^1^, MINE_enhanced Hi-C can hit more significant interactions than the raw high-resolution Hi-C.

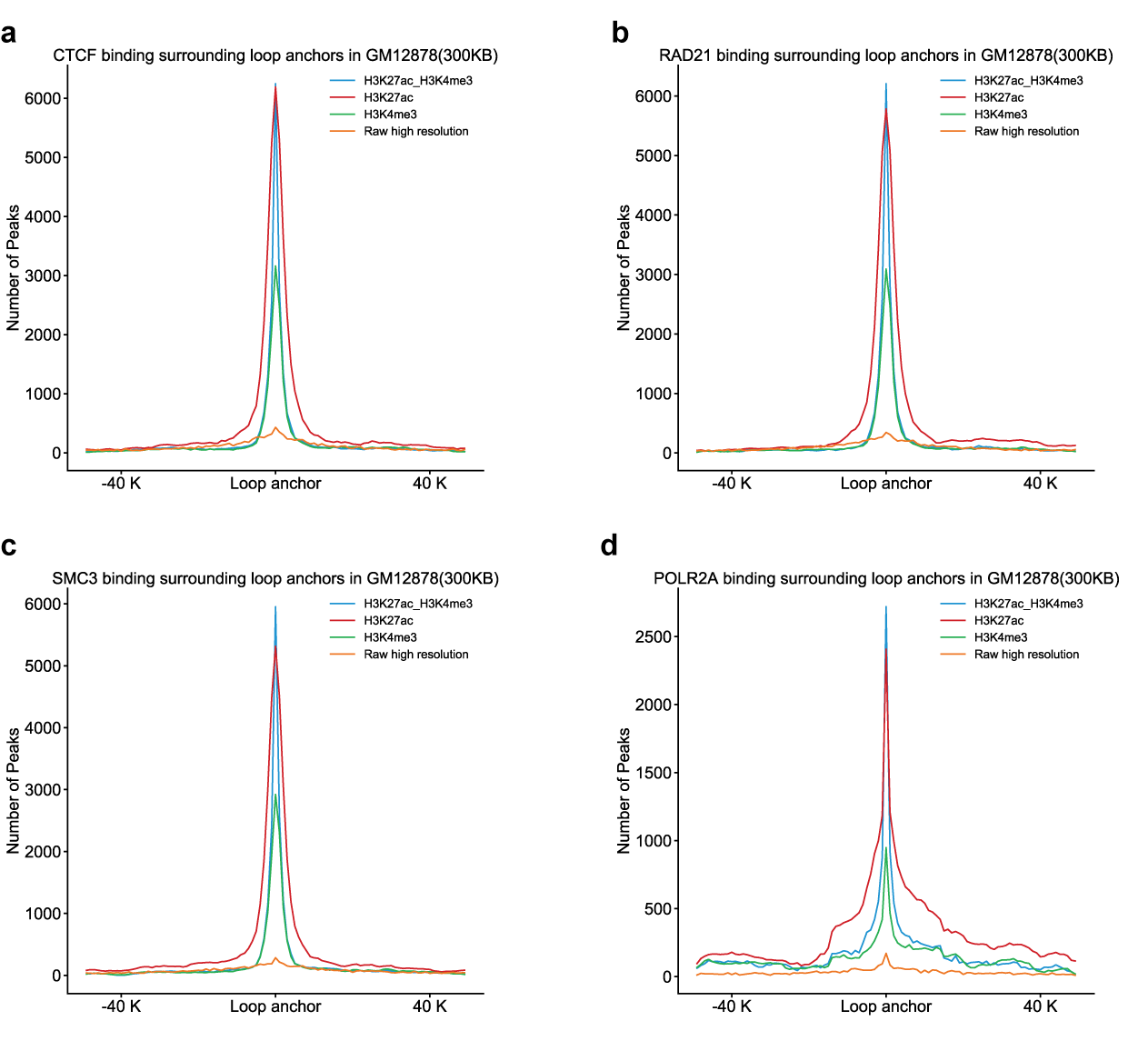

**Figure S4-** the comparison of results obtained by training in different combinations within a genome distance of 2-300kb.

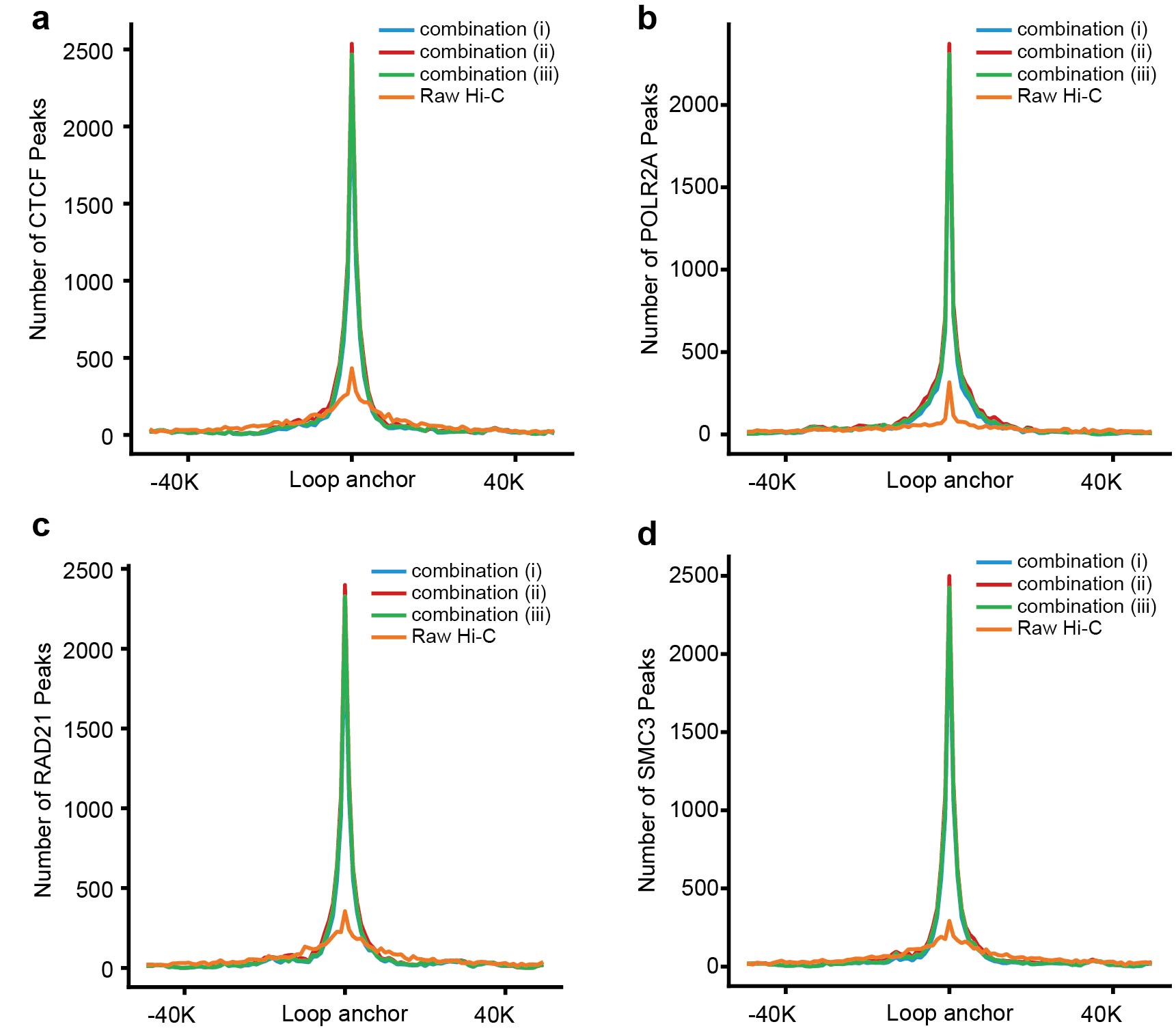

**Figure S5.** Predictions for IMR90 cell line (raw reads:1.08 billion) using the model trained in GM12878 cell line with ATAC-seq, H3K27ac ChIP-seq, and H3K4me3 ChIP-seq data. The following three combinations are chosen as input to predict. Combination (i): ATAC-seq, H3K27ac ChIP-seq, H3K4me3 ChIP-seq (i.e., blue lines). Combination (ii): ATAC-seq, H3K27ac ChIP-seq (i.e., red lines). Combination (iii): ATAC-seq, H3K4me3 ChIP-seq (i.e., green lines). a-d. The loops within the range of 2-100kb genome distance called from active model using different combinations as input anchoring to the CTCF, RAD21, SMC3, POLR2A factors.

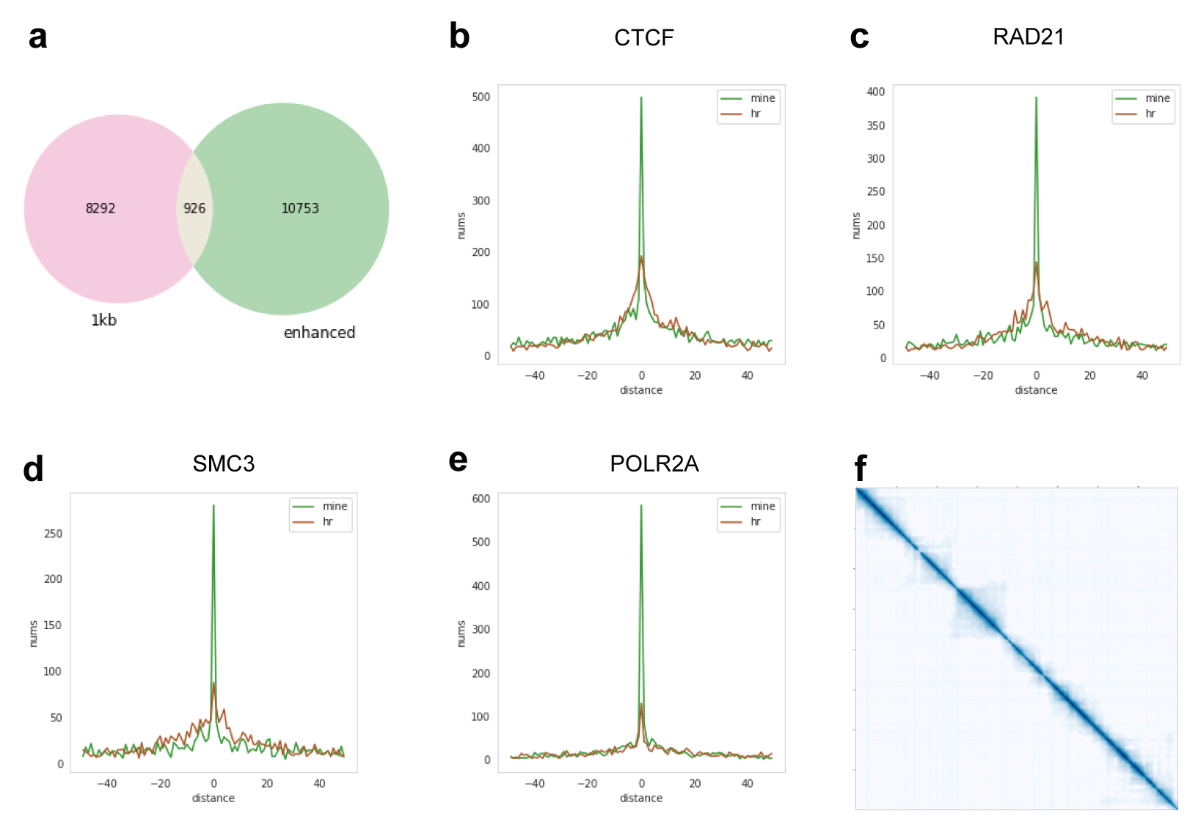

**Figure S6** the prediction result using the active model trained with ATAC-seq, H3k27ac, H3K4me3 ChIP-seq of GM12878 cell line. We use the ATAC-seq, H3K27ac ChIP-seq and H3K4me3 ChIP-seq as the input of prediction for **K562** cell line data (reads number:0.9 billion). (a) the overlap of loops detected from raw high-resolution Hi-C and MINE_enhanced Hi-C by FitHi-C2. (b-e) The hit distance distribution graph for the biologically meaningful interaction (CTCF, rad21, SMC3, POLR2A) within the 2-100kb genome distance. (f) the heat map of the MINE_enhanced Hi-C.

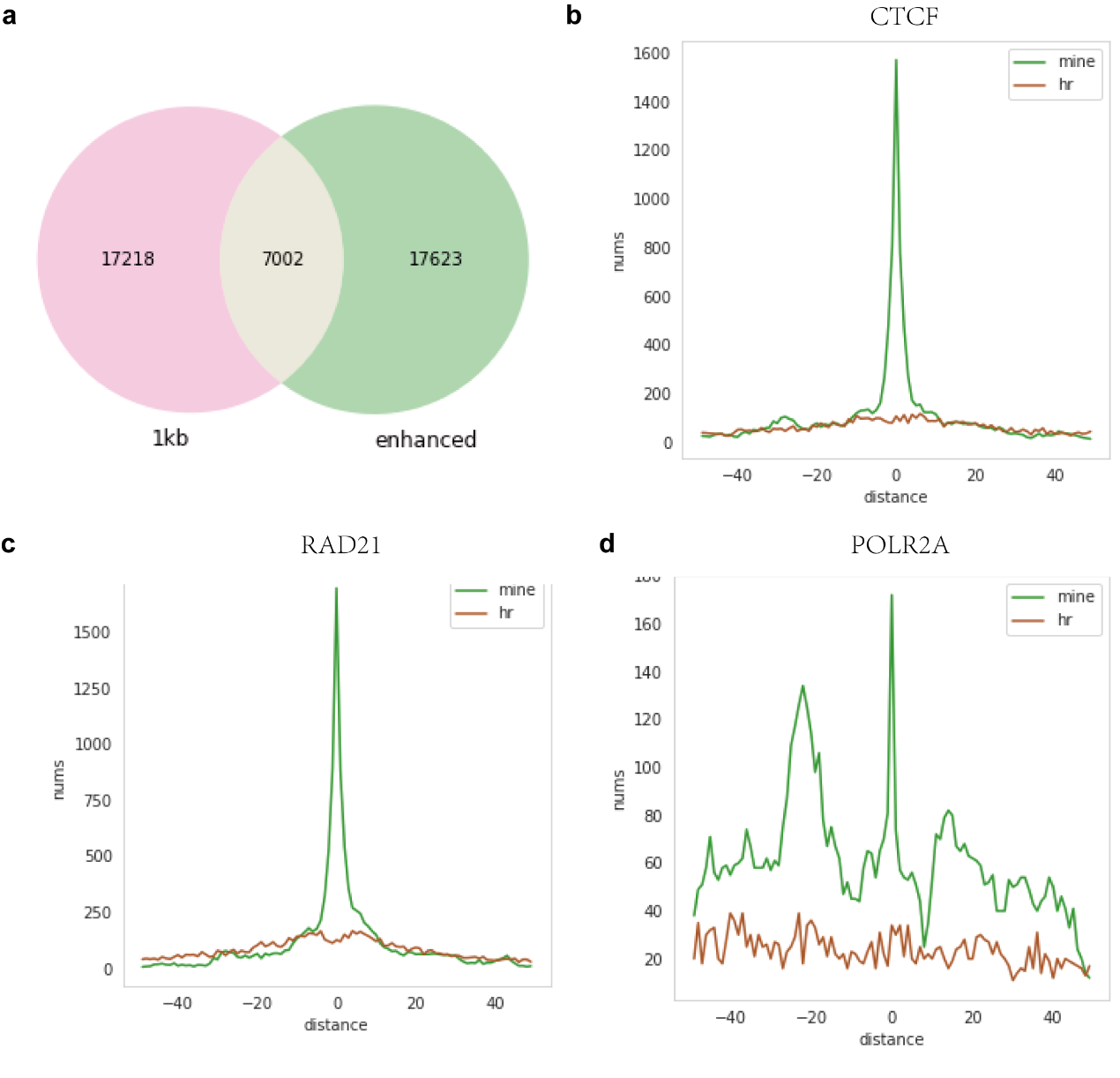

**Figure S7** the prediction result using the active model trained with ATAC-seq, H3k27ac, H3K4me3 ChIP-seq of GM12878 cell line. We use the ATAC-seq, H3K27ac ChIP-seq and H3K4me3 ChIP-seq as the input of prediction for **H1** cell line data (reads number:3.22 billion). (a) the overlap of loops detected from raw high-resolution Hi-C and MINE_enhanced Hi-C by FitHi-C2. (b-d) The hit distance distribution graph for the biologically meaningful interaction (CTCF, rad21, POLR2A) within the 2-100kb genome distance.

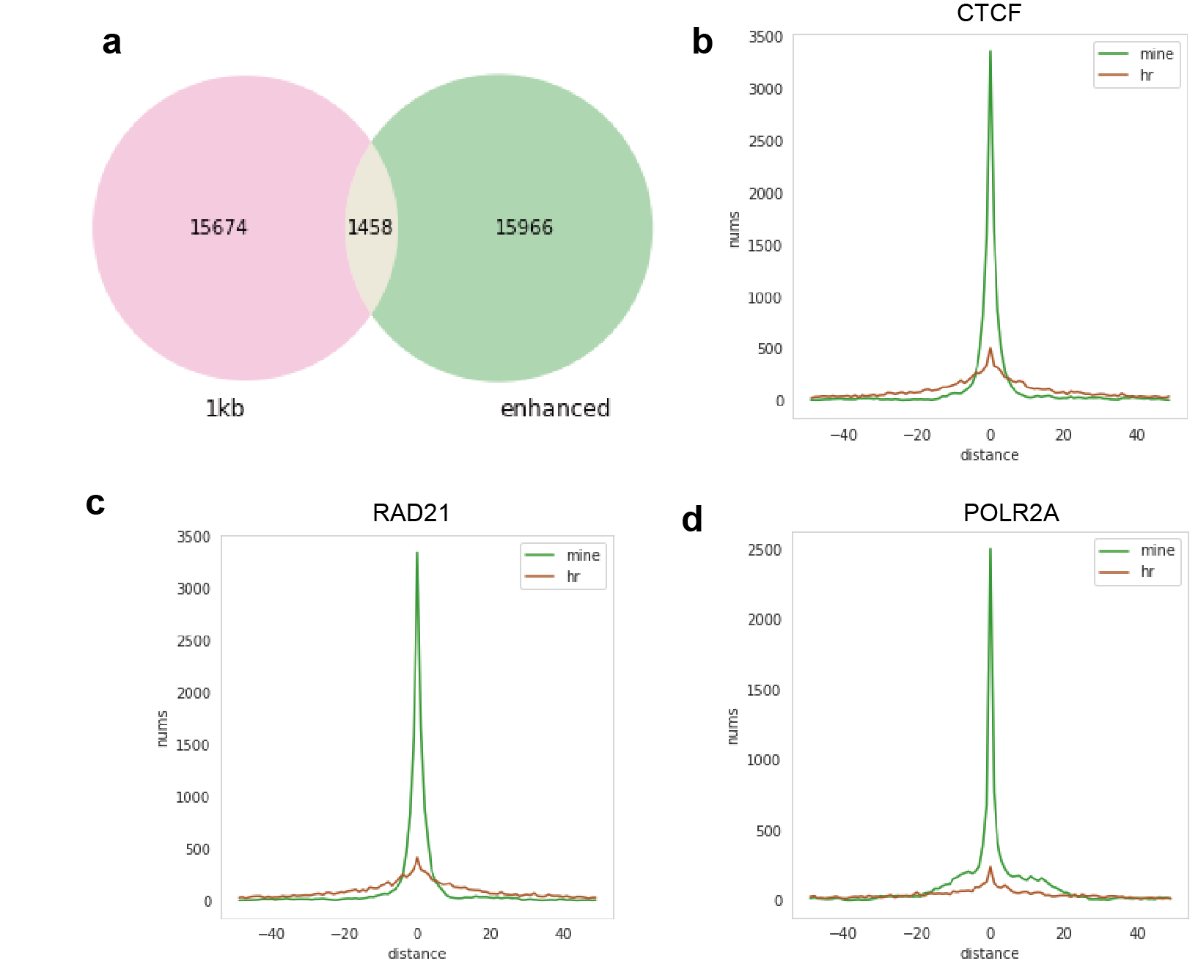

**Figure S8-** the prediction result using the active model trained with ATAC-seq, H3k27ac, H3K4me3 ChIP-seq of GM12878 cell line. We use the ATAC-seq, H3K27ac ChIP-seq and H3K4me3 ChIP-seq as the input of prediction for **HepG2** cell line data (reads number:2 billion). (a) the overlap of loops detected from raw high-resolution Hi-C and MINE_enhanced Hi-C by FitHi-C2. (b-d) The hit distance distribution graph for the biologically meaningful interaction (CTCF, rad21, POLR2A) within the 2-100kb genome distance.

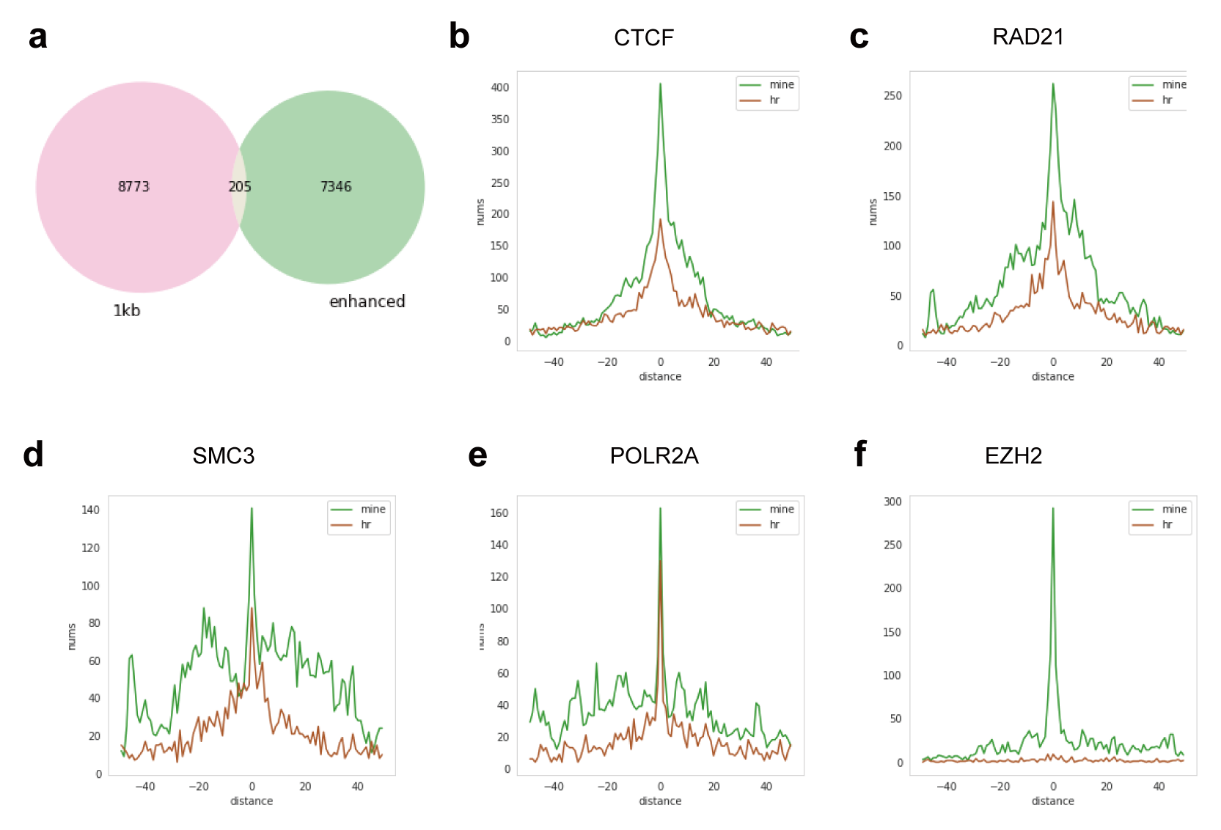

**Figure S9-**the repressive model can enhance the low-resolution Hi-C of **K562** cell line using the GM12878 cell line data to train the model.

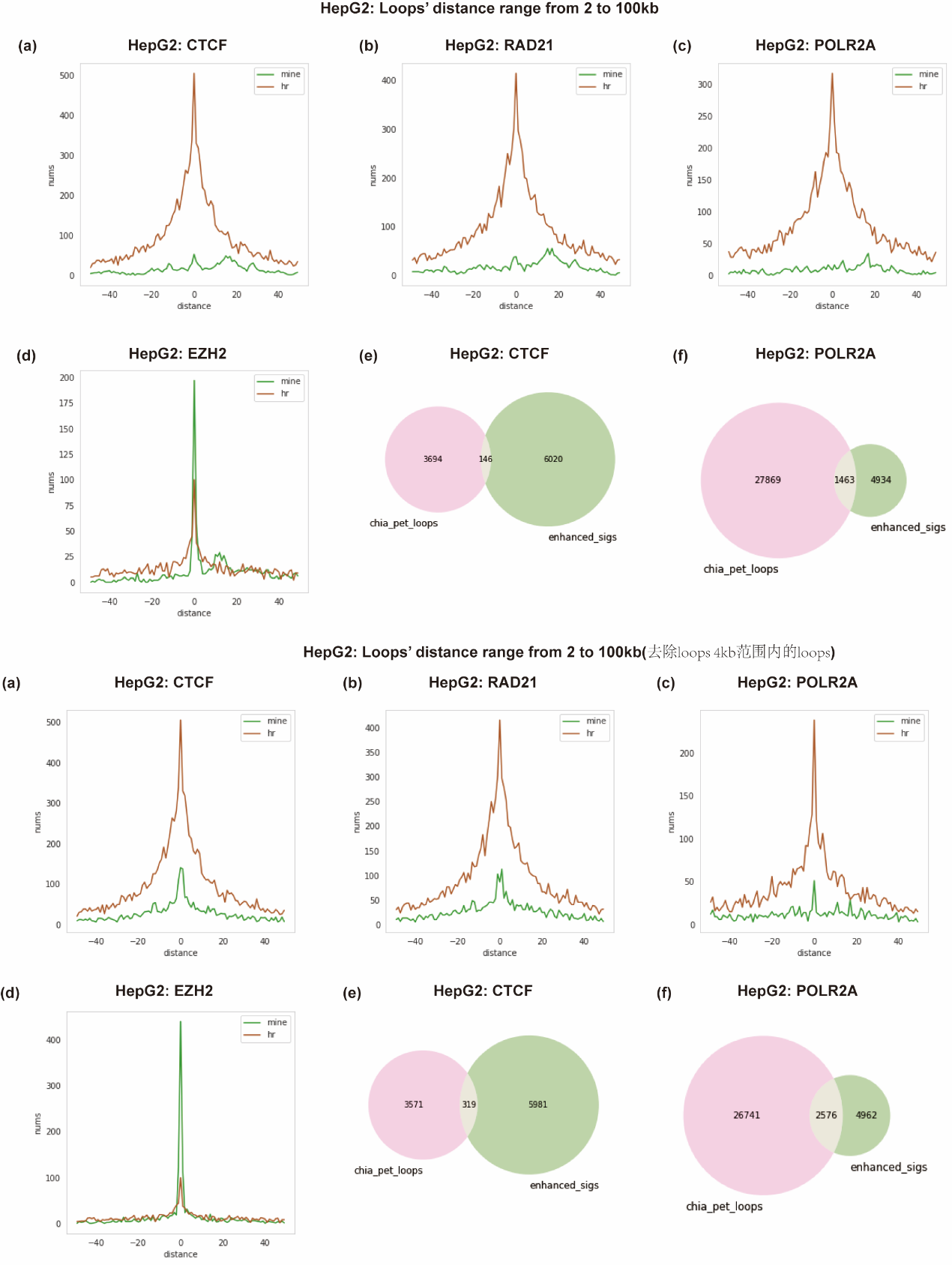

**Figure S10-**the repressive model can enhance the low-resolution Hi-C of **HepG2** cell line using the GM12878 cell line data to train the model. (a-d) The hit distance distribution graph for the biologically meaningful interaction (CTCF, rad21, POLR2A) within the 2-100kb genome distance. (e-f) The overlap of the loops detected by FitHi-C2 that hit the CTCF, POLR2A factors and the corresponding Chia-PET interactions.

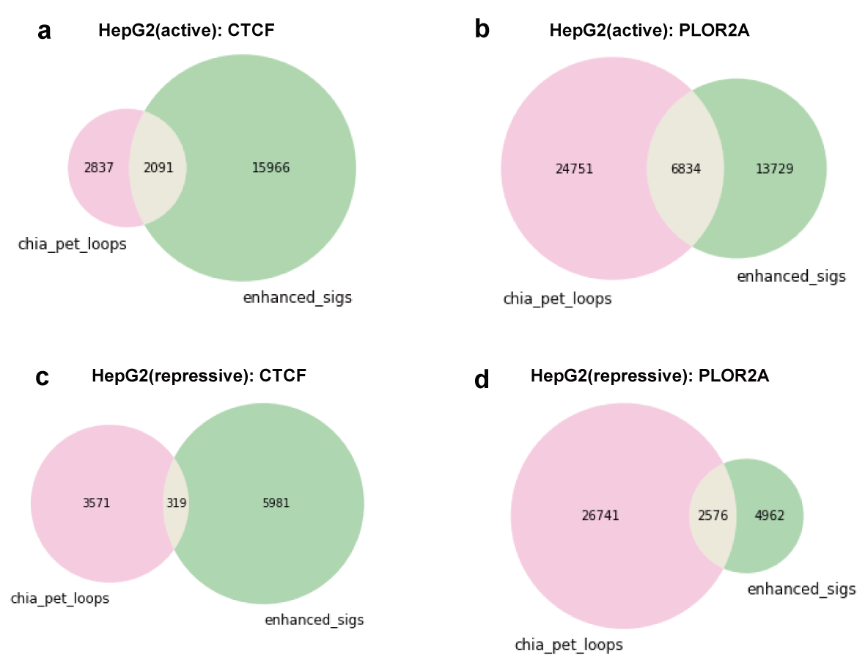

**Figure S11 –**The MINE-enhanced Hi-C can overlap the ChIA-PET data of related factors at short distances. The active and repressive models trained by GM12878 were used to enhance the Hi-C data of **HepG2** cells, and the FitHi-C2 was used to predict the loops. The overlap of the loops that hit the CTCF, POLR2A factors and the corresponding ChIA-PET interactions.

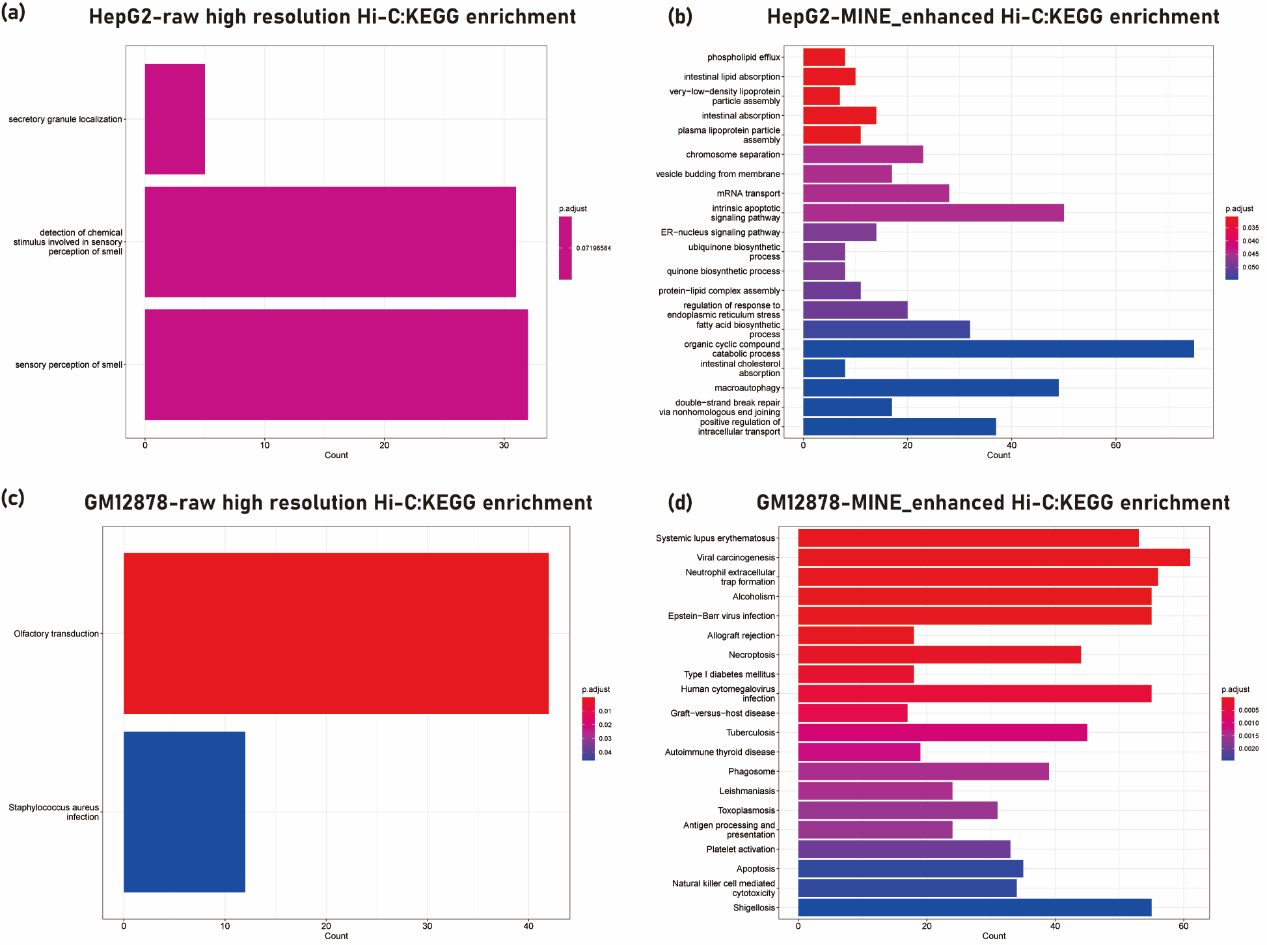

**Figure S12 –**The gene KEGG pathway enrichment results from MINE and raw high-resolution Hi-C’s specific transcription start site of **HepG2** cell line (loops are detected by FitHi-C2^1^).
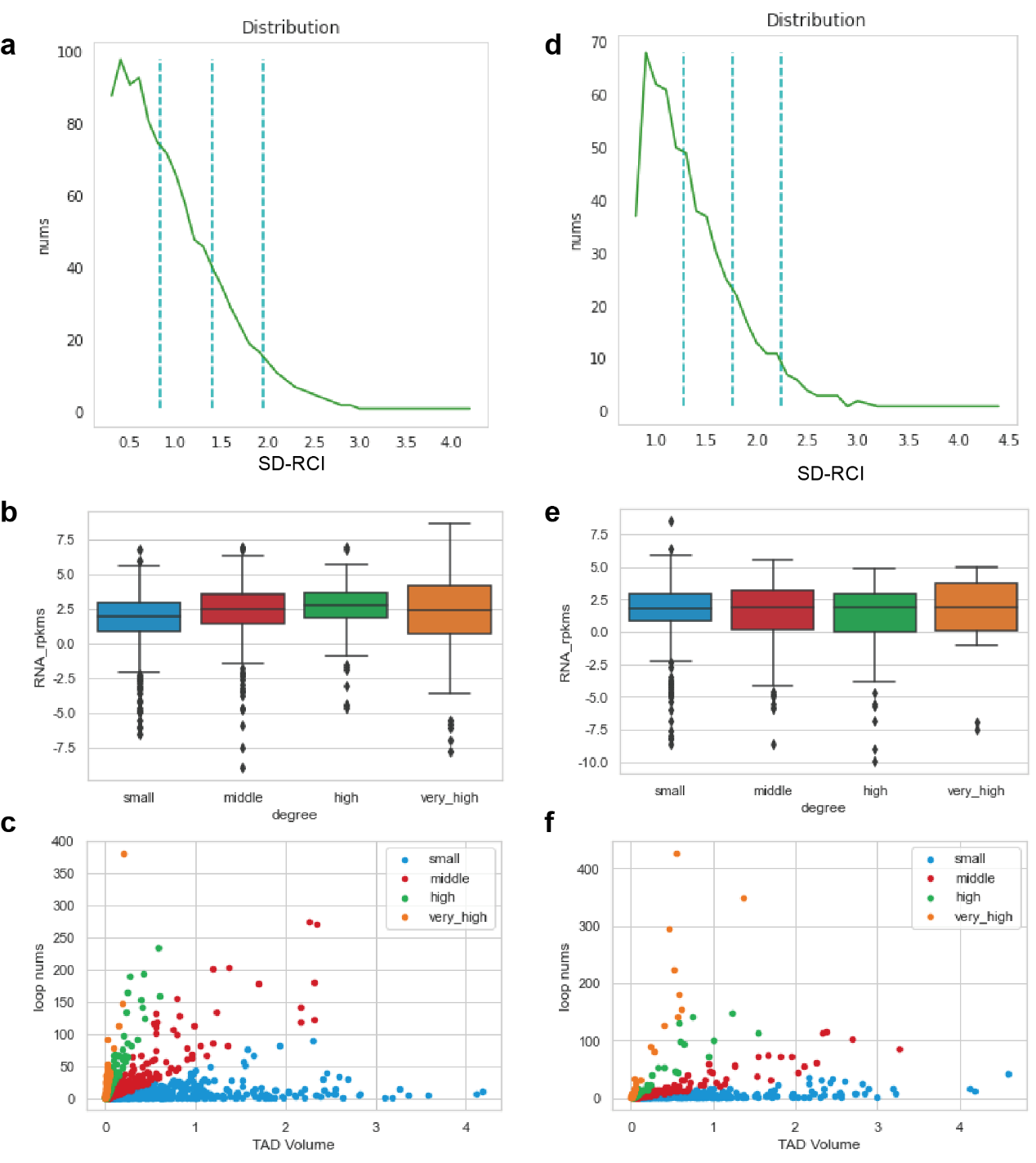

**Figure S13-**Active model could get a higher SD-RCI. (a, b, c) are calculated at the genomic distance of 2-100kb, (d, e, f) are calculated at the genomic distance of 2-300kb. (a) the distribution of the number of active loops change with the SD-RCI value calculated by SD-RCI method. (b) the box plot of RPKM and SD-RCI degree, we divide the SD-RCI into four degrees according to the value of δ in the Gaussian distribution of figure (a). (c) the dot plot of the number of loops and volume of TADs.

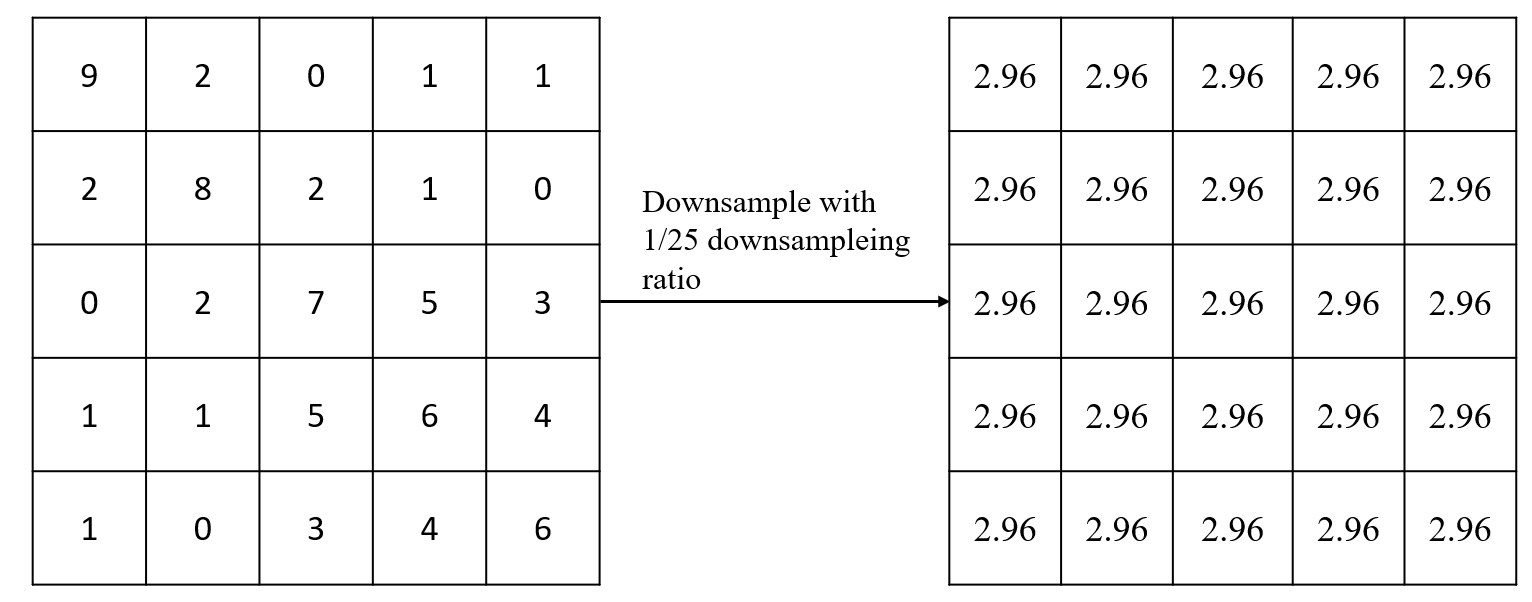

**Figure S14-**Example of downsampling Hi-C data with 1/25 downsampling ratio.

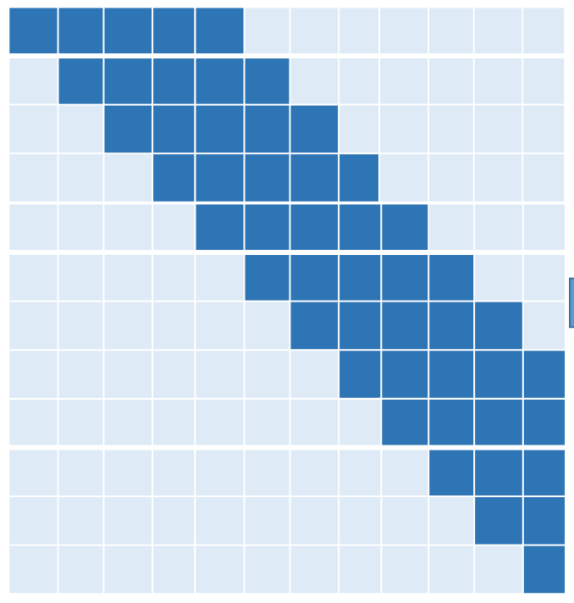

**Figure S15-**Schematic diagram of matrix-based sample division.

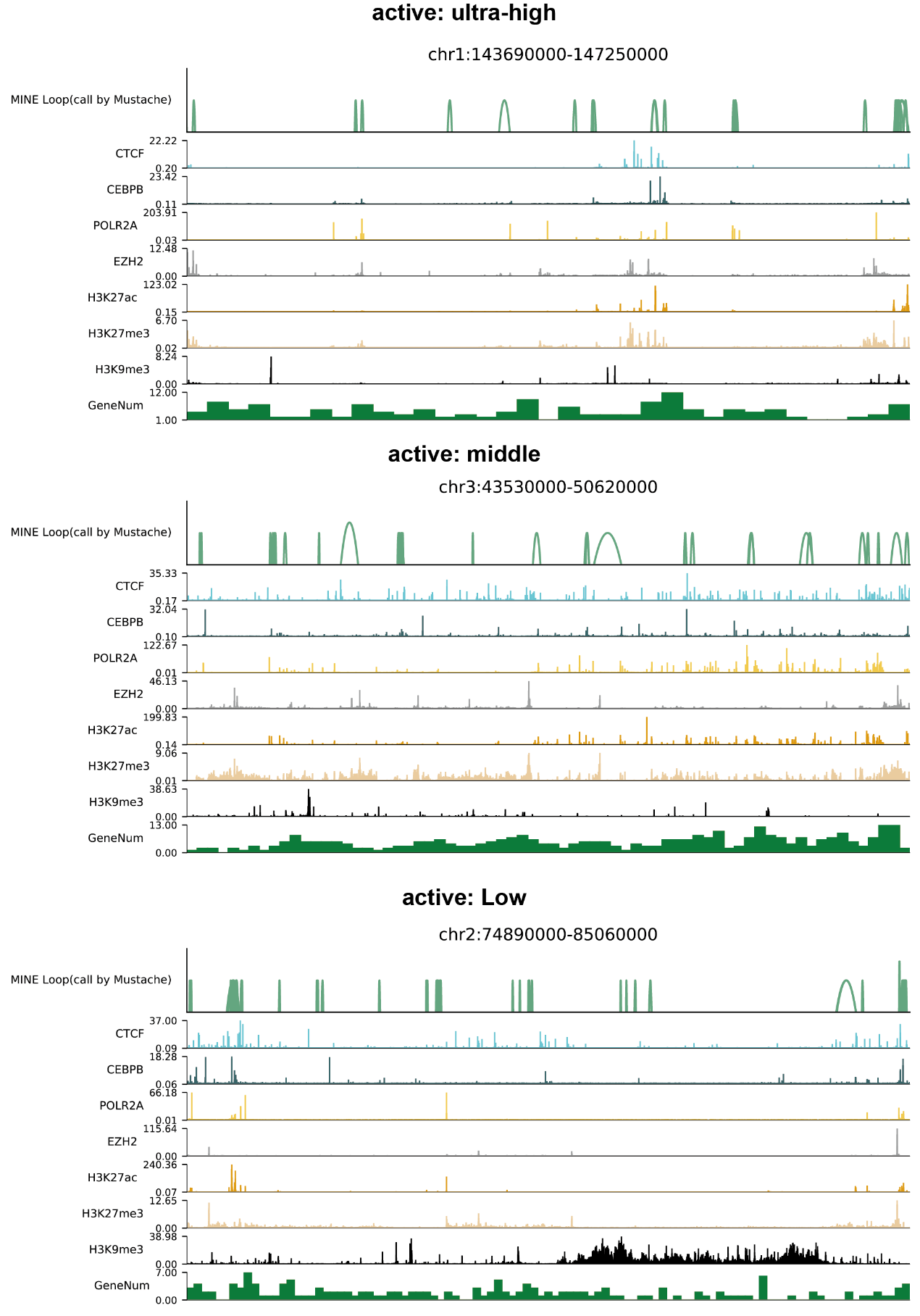

**Figure S16** The visualization of loops and CTCF, histone mark tracks for ultra-high, middle, low SD-RCI level in active state.

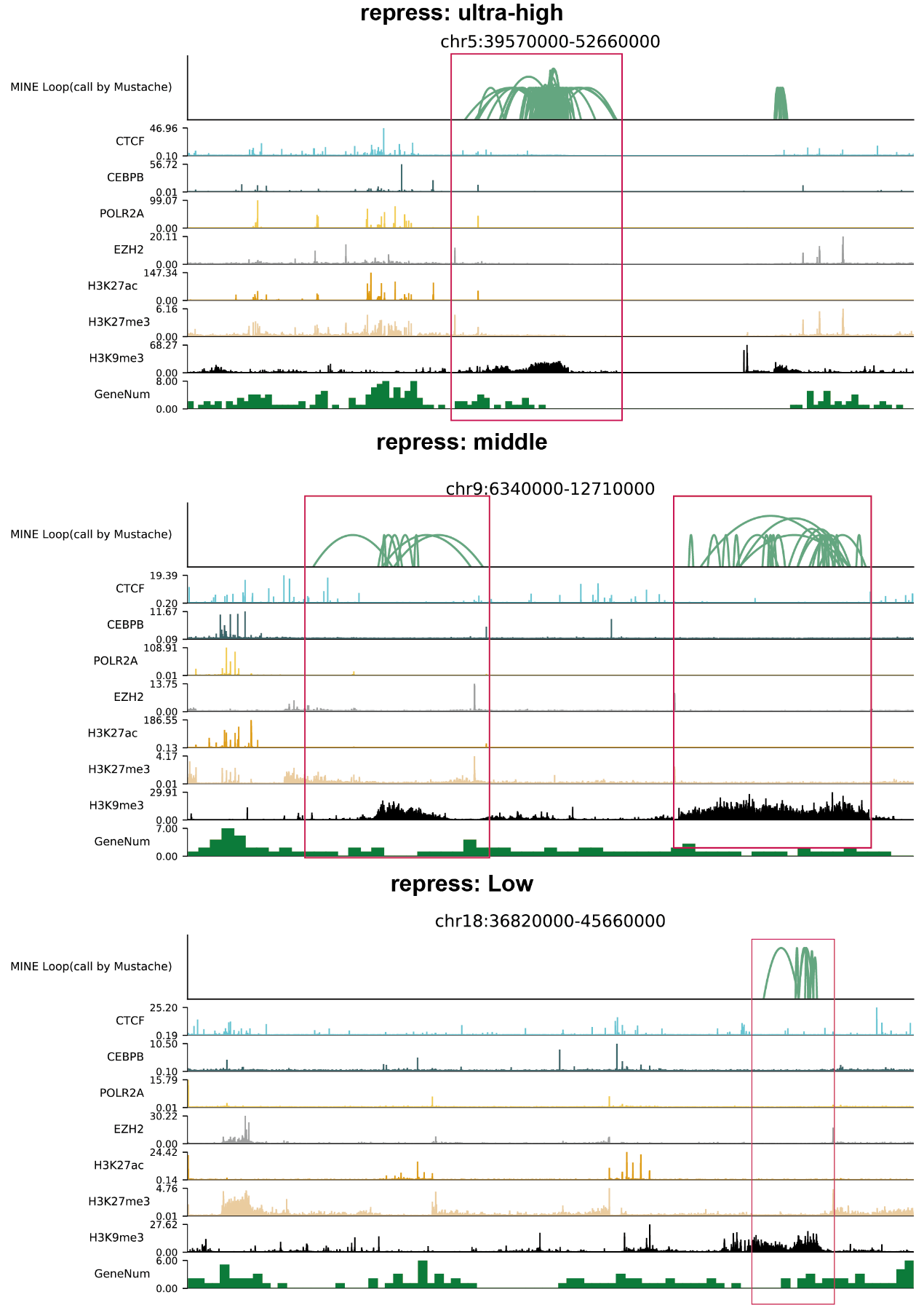

**Figure S17** The visualization of loops and CTCF, histone mark tracks for ultra-high, middle, low SD-RCI level in repress state.

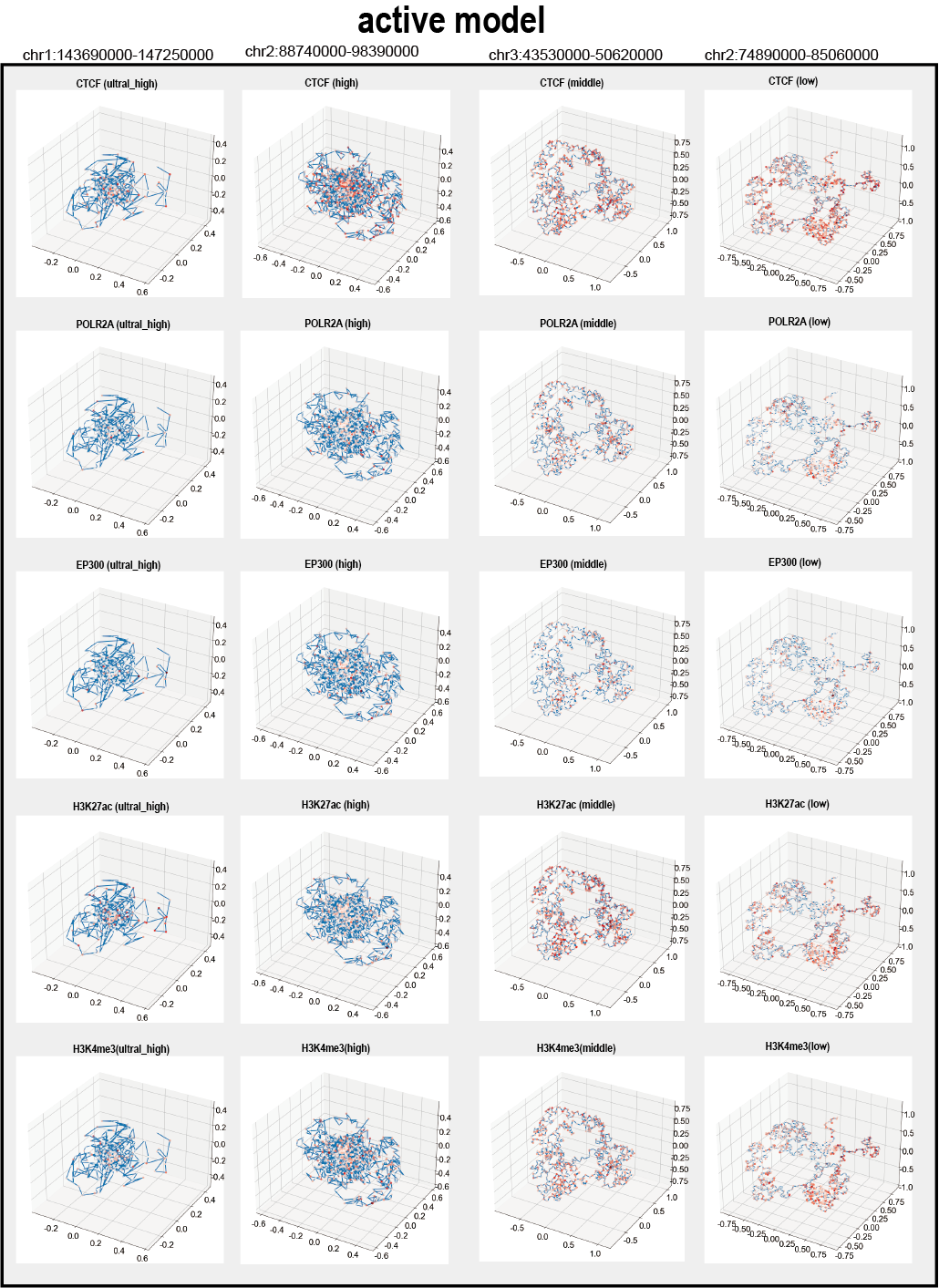

**Figure S18-** The distribution of various transcription factors in the three-dimensional space of chromatin corresponding to the active models.

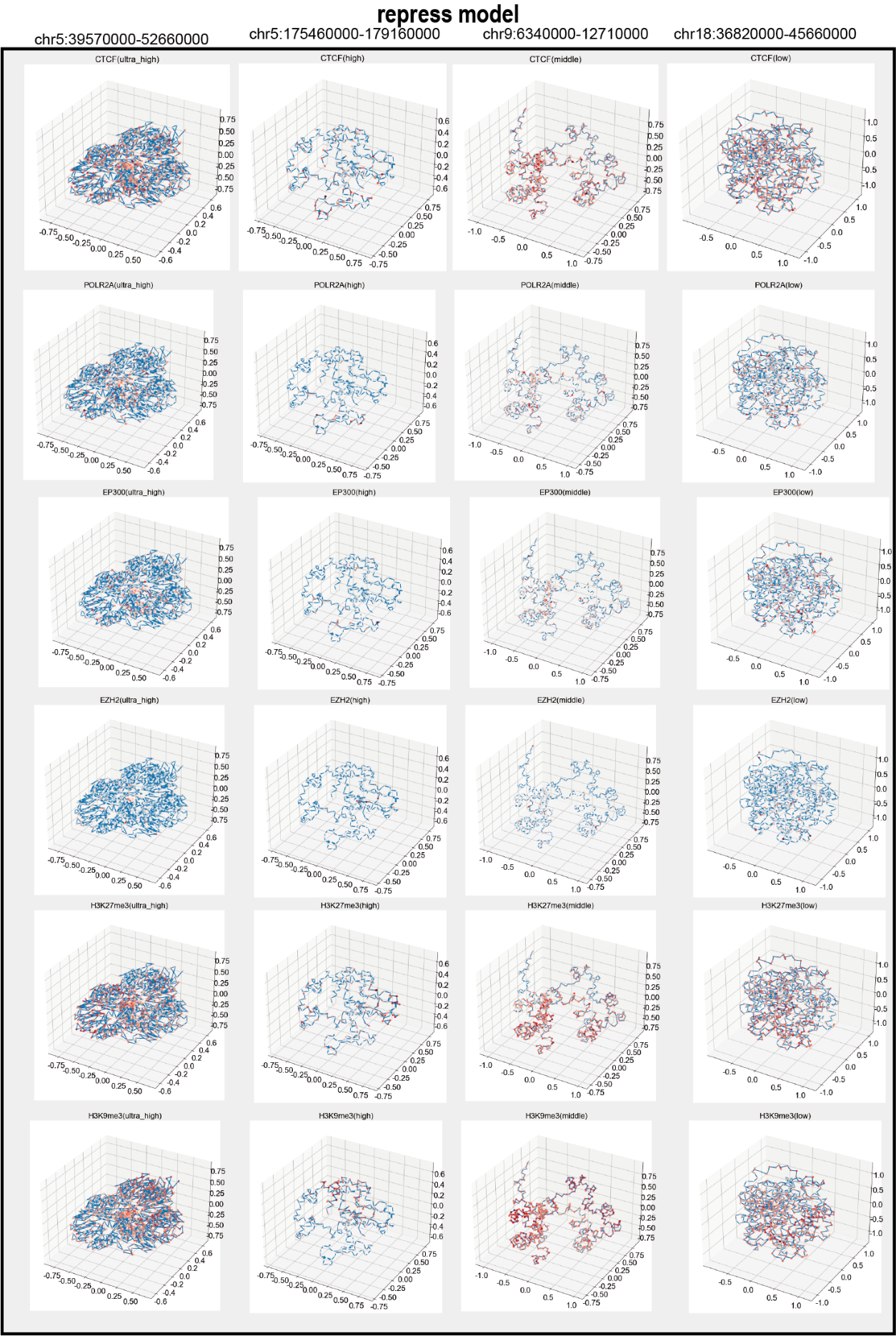

**Figure S19-** The distribution of various transcription factors in the three-dimensional space of chromatin corresponding to the repress models.

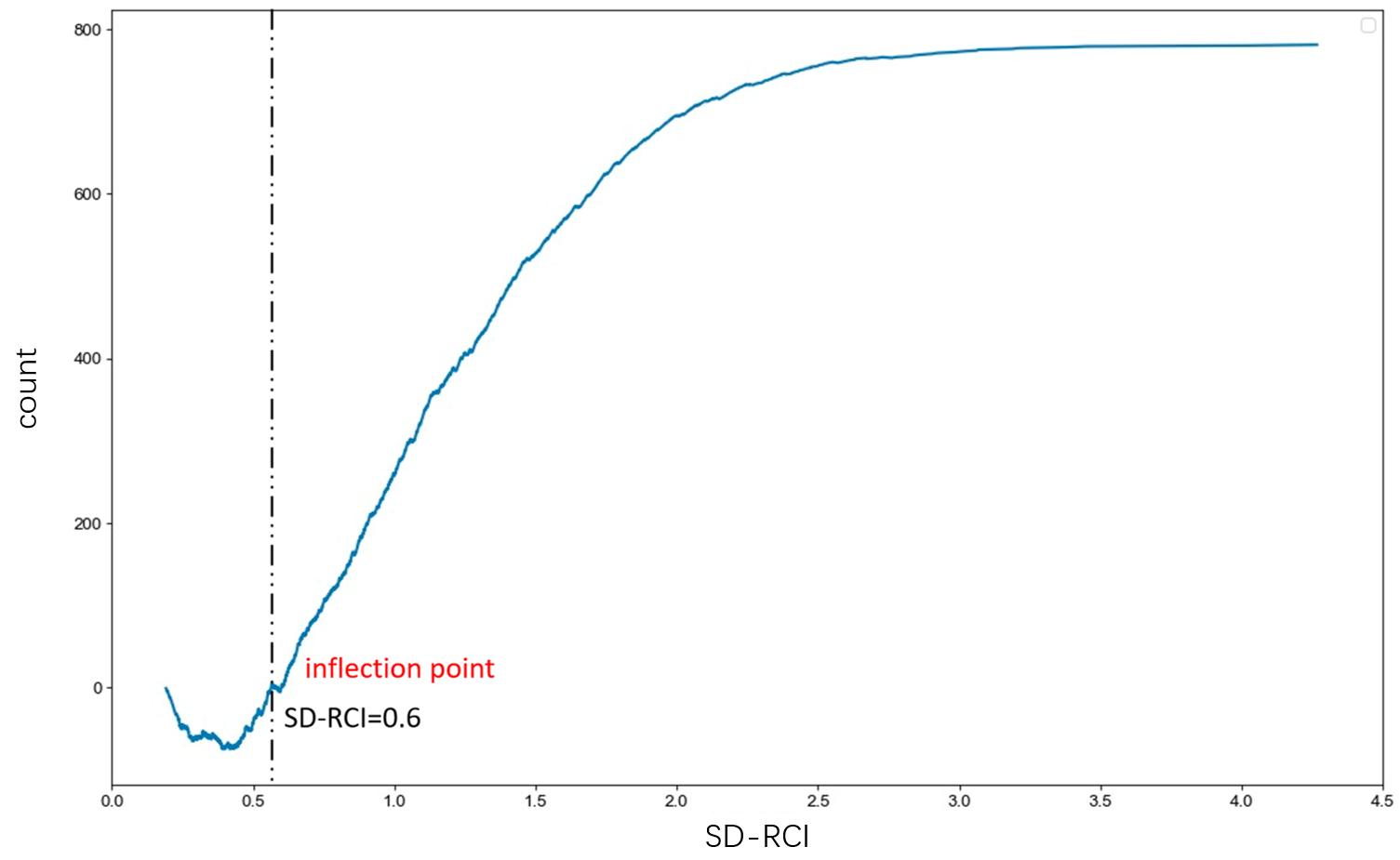

**Figure S20-The curve of count and SD-RCI, where the count is calculated as follows: when the TAD volume increases, count+1, then count-1.**

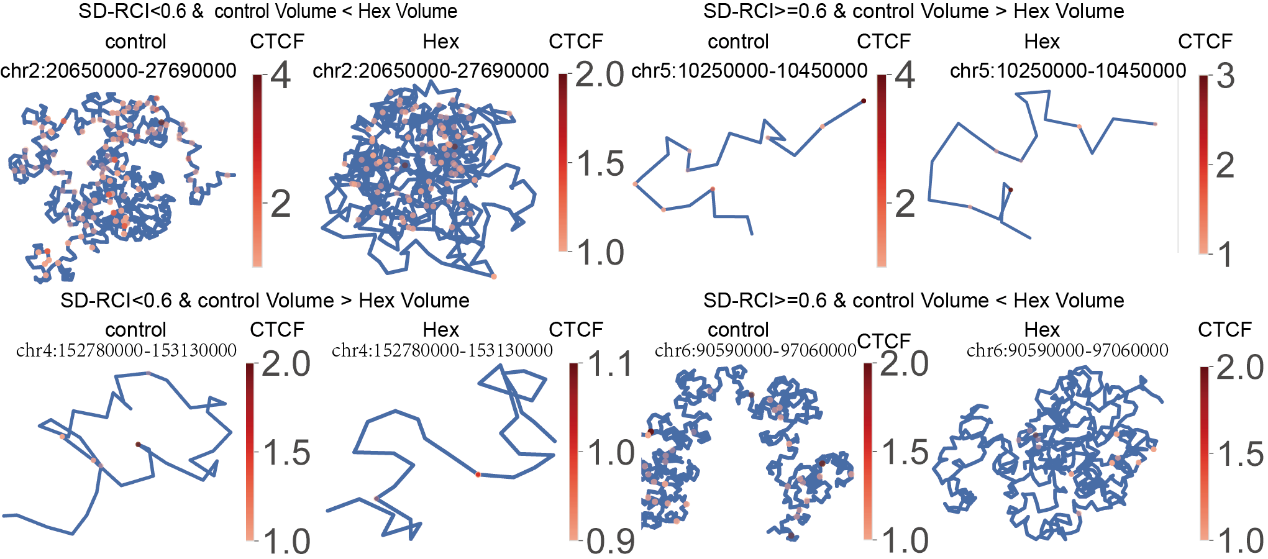

**Figure S21-** The 3D structure visualization marked with CTCF anchor intensity in four types of TADs in control group and Hex group.

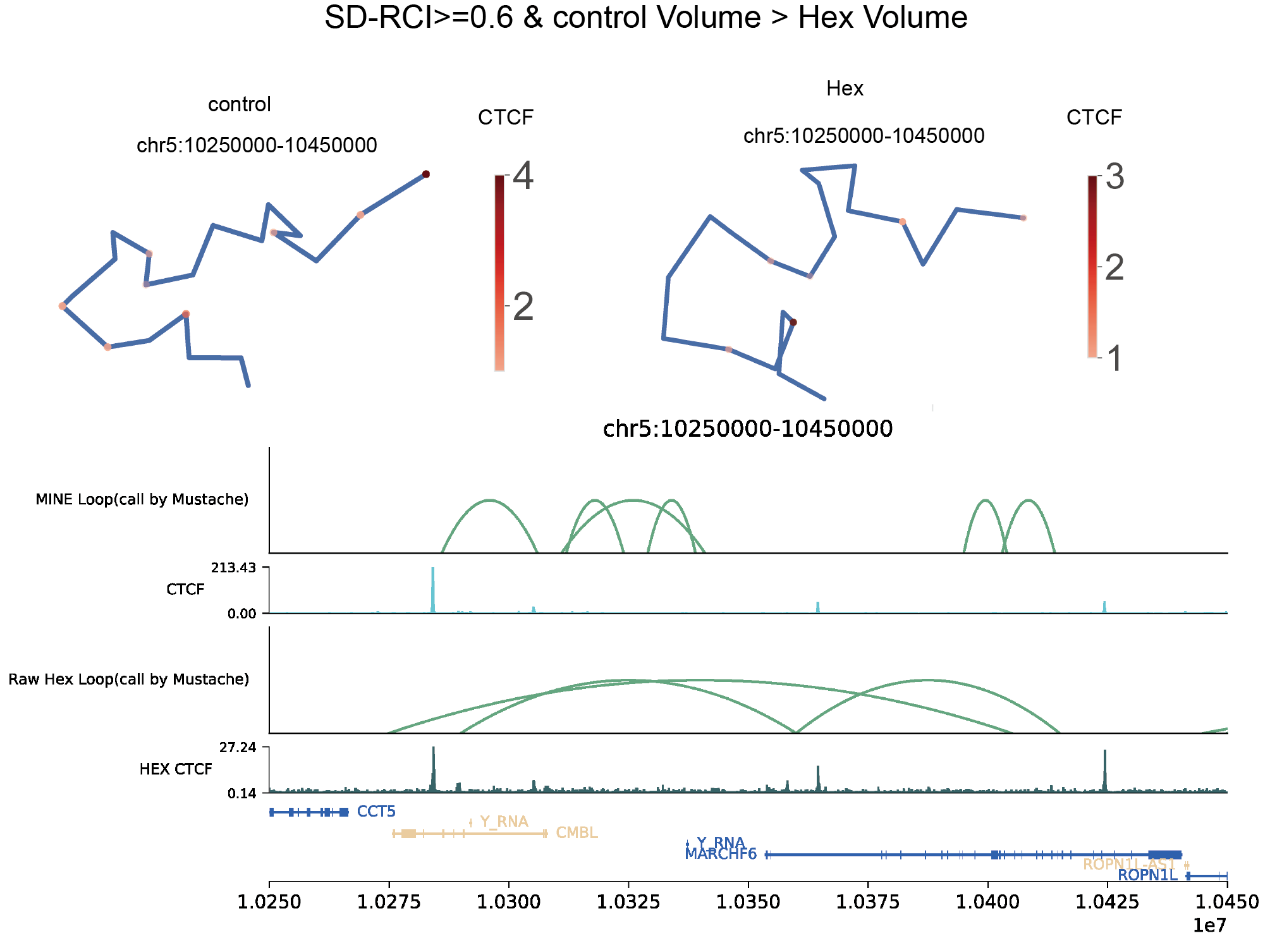

**Figure S22-**a. The 3D structure marked with CTCF anchor intensity of TAD typed with “SD-RCI>=0.6 & control Volume > Hex Volume”. c. The visualization of loops and tracks from control group and Hex group.

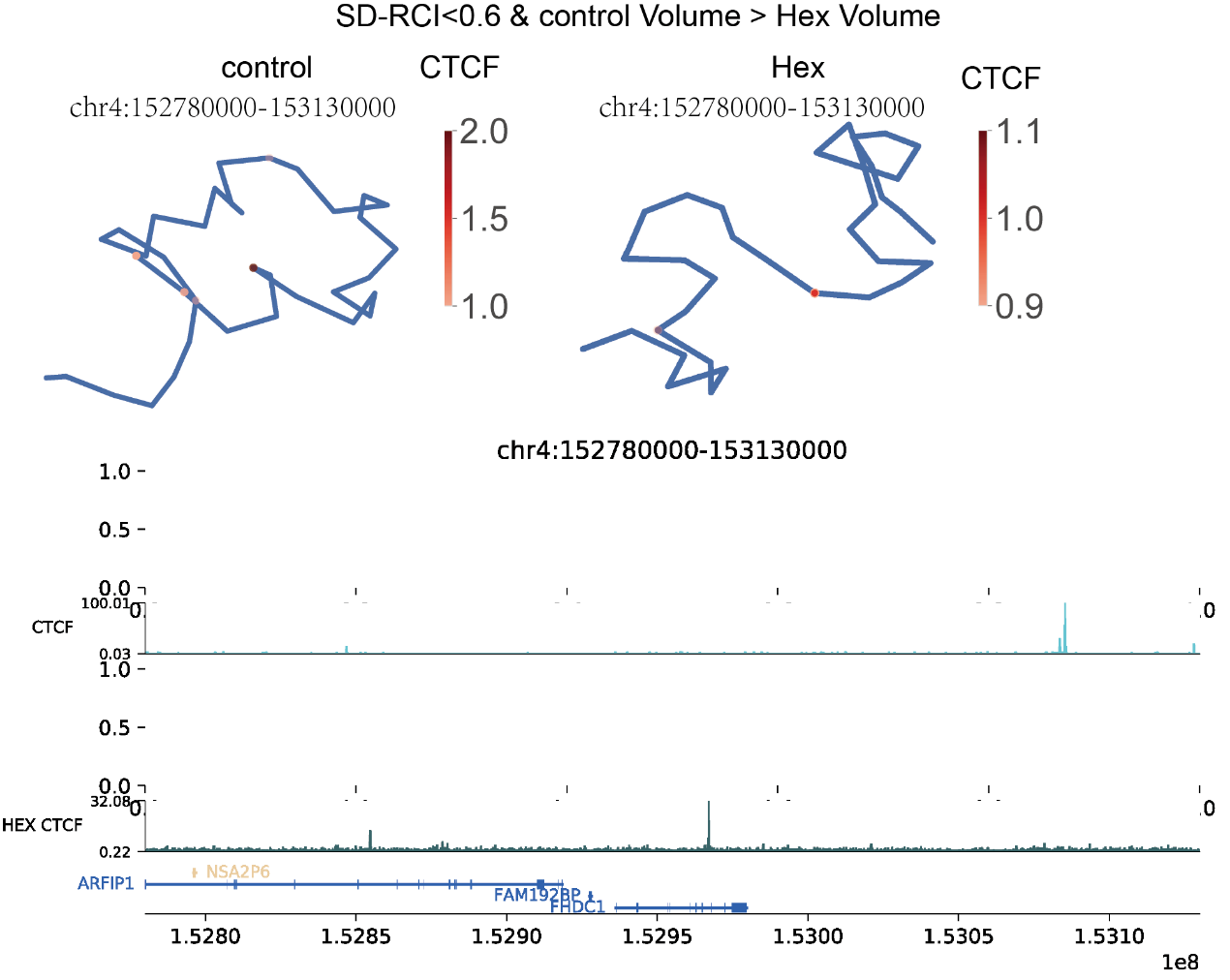

**Figure S23-**a. The 3D structure marked with CTCF anchor intensity of TAD typed with “SD-RCI<0.6 & control Volume > Hex Volume”. c. The visualization of loops and tracks from control group and Hex group.

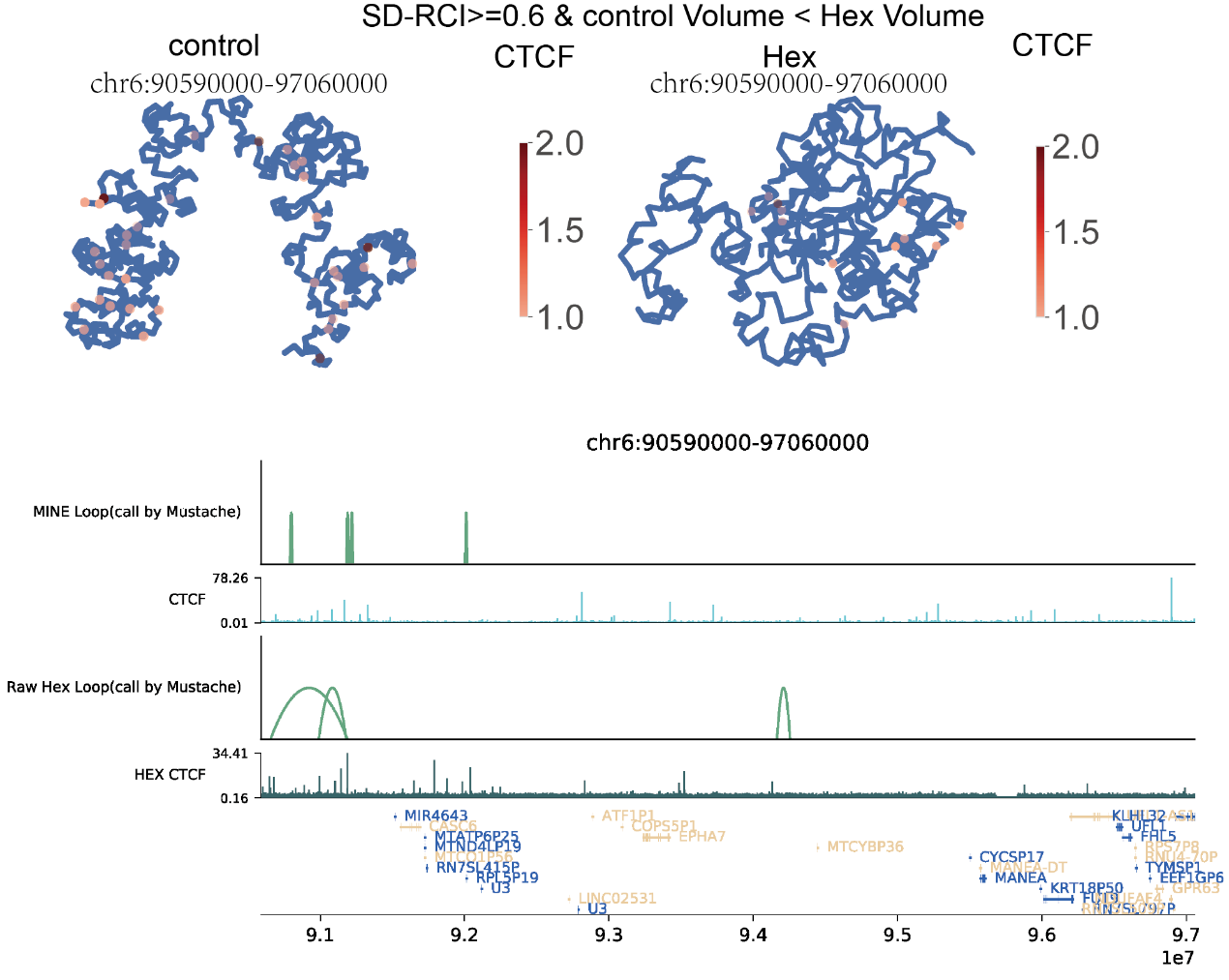

**Figure S24-**a. The 3D structure marked with CTCF anchor intensity of TAD typed with “SD-RCI>=0.6 & control Volume < Hex Volume”. c. The visualization of loops and tracks from control group and Hex group.
